# Overcoming the Blood-Brain Barrier for Gene Therapy via Systemic Administration of GSH-Responsive Silica Nanocapsules

**DOI:** 10.1101/2022.10.27.513950

**Authors:** Yuyuan Wang, Xiuxiu Wang, Ruosen Xie, Jacobus C. Burger, Yao Tong, Shaoqin Gong

**Affiliations:** Department of Ophthalmology and Visual Sciences, University of Wisconsin-Madison, Madison, WI 53705, USA; Department of Biomedical Engineering, University of Wisconsin-Madison, Madison, WI 53706, USA; Wisconsin Institute for Discovery, University of Wisconsin-Madison, Madison, WI 53715, USA; Department of Materials Science and Engineering, University of Wisconsin-Madison, Madison, WI 53706, USA; Department of Chemistry, University of Wisconsin-Madison, Madison, WI 53706, USA

**Author notes:** Correspondence should be addressed to SG.

## Abstract

CRISPR genome editing has demonstrated great potential to treat the root causes of many genetic diseases, including central nervous system (CNS) disorders. However, the promise of brain-targeted therapeutic genome editing relies on the efficient delivery of biologics bypassing the blood-brain barrier (BBB), which represents a substantial challenge in the development of CRISPR therapeutics. In this study, we created a library of GSH-responsive silica nanocapsules (SNCs) and screened them for brain targeting via systemic delivery of nucleic acids and CRISPR genome editors. *In vivo* studies demonstrated that systemically delivered SNCs conjugated with glucose and RVG peptide under glycemic control can efficiently bypass the intact BBB, enabling brain-wide delivery of various biologics (mRNA, Cas9 mRNA/sgRNA, and Cas9/sgRNA ribonucleoprotein) targeting both exogenous genes (i.e., Ai14 stop cassette) and disease-relevant endogenous genes (i.e., *App* and *Th* genes) in Ai14 reporter mice and wild-type mice, respectively. In particular, we observed up to 28% neuron editing via systemic delivery of Cre mRNA in Ai14 mice, up to 6.1% amyloid precursor protein (*App*) gene editing (resulting in 19.1% reduction in the expression level of intact APP), and up to 3.9% tyrosine hydroxylase (*Th*) gene editing (resulting in 30.3% reduction in the expression level of TH) in wild-type mice. This versatile SNC nanoplatform may offer a novel strategy for the treatment of CNS disorders including Alzheimer’s, Parkinson’s, and Huntington’s disease.

## 1. Introduction

Clustered regularly interspaced short palindromic repeats (CRISPR) genome editing is a revolutionary and versatile genome editing technique with wide-ranging utility^[1]^. The CRISPR/Cas9 system can target and edit disease-causing mutations in a sequence-dependent manner, resulting in a permanent genetic change and thus offering the promise to cure genetic diseases ^[1]^. *In vivo* somatic genome editing using CRISPR/Cas9 is anticipated to be the next wave of therapeutics for many major health threats, including central nervous system (CNS) disorders (e.g., Alzheimer’s disease, Parkinson’s disease and Huntington’s disease) ^[2]^. However, the promise of brain gene therapy relies on the efficient delivery of biologics (e.g., nucleic acids and CRISPR genome editor) to the brain, which is extremely challenging due to the blood-brain barrier (BBB) ^[3]^.

To date, *in vivo* brain genome editing has been mostly mediated by viral vectors that require laborious customization for every target, have limited DNA packaging capacity, and also have poor biosafety profiles, including immunogenicity and rare but dangerous integration events ^[4]^. Production scale-up of viral vectors is also difficult. While non-viral nanocarriers have been explored for brain gene therapy, these nanocarriers are mostly administered via intracranial administration, which is invasive and only suitable for gene therapy in a small and localized brain region around the injection site ^[5]^. There are several recent reports of intravenously injected nanoparticles for the delivery of small proteins (e.g., Cre recombinase) ^[6]^ or siRNA/antisense oligonucleotide (ASO) ^[7]^ into the brains of healthy mice or diseased mice. There are also a couple of recent reports of intravenous delivery of large biologics such as CRISPR plasmid and Cas9 protein/sgRNA ribonucleoprotein (RNP) bypassing the BBB; however, these studies were carried out in disease models (e.g., brain tumor or Alzheimer’s disease models) with compromised BBB integrity [8]. For instance, the Cas9 RNP delivered by an RNP nanocage using Angiopep-2 peptide as a targeting ligand showed more than 20-fold higher accumulation in brain tumors than normal brain tissue ^[8b]^. Thus far, there are no reports on systemic delivery of mRNA, Cas9 mRNA/sgRNA, and Cas9 RNP targeting the whole brain in healthy mice with intact BBB. Additionally, prior studies often lack information on types of brain cells edited and their respective editing efficiency as well as the *in vivo* biodistribution (i.e., %ID/g, percent injected dose per gram tissue), substantiating an urgent need to develop safe and efficient non-viral delivery vehicles capable of systemically delivering a wide range of biologics for whole-brain therapeutic genome editing.

A number of peptide- or protein-based targeting ligands were employed to enhance the ability of nanocarriers to bypass the BBB including rabies virus glycoprotein (RVG) peptide, transferrin, and apolipoprotein E targeting nicotinic acetylcholine receptor (nAChR), transferrin receptor, and low-density lipoprotein receptor, respectively ^[9]^. RVG peptide, derived from rabies virus glycoprotein, was known to specifically interact with nAChR on neuronal cells and brain capillary endothelial cells (BCECs), thereby facilitating the RVG-conjugated NPs to cross the BBB and enhancing their neuronal uptake ^[10]^. However, these targeting strategies only led to modest success in delivering nanocarriers to the brain (e.g., less than 3% dose/g brain tissue, more commonly less than 1% dose/g). Recently, Kataoka et al. reported a new strategy to bypass the BBB via glucose transporter-1 (GLUT1) mediated transcytosis through glycemic control ^[7b, 11]^. Nanoparticles conjugated with glucose were able to achieve 5-7% dose/g in brain tissue ^[7b, 11b]^. GLUT1 is expressed on the BCECs. Upon 16-24 h fasting, the density of GLUT1 on the luminal side of the cell membrane is increased. When the blood glucose is rapidly restored, GLUT1 on the luminal side starts to migrate to the abluminal side of the BCEC membrane, presumably through a transcytosis process (Fig. 1C) ^[11b, 12]^. Glucose-conjugated nanocarriers can bypass the BBB utilizing this mechanism. However, bypassing the BBB is only the first step for nanoparticle-mediated genome editing in the brain. Efficient delivery of genome editors into brain cells involves multiple steps including but not limited to: diffusion in the brain parenchyma, recognition by target cell(s), endocytosis, endo/lysosomal escape, and payload release in the cytosol. Thus, achieving high genome editing efficiency in the brain requires both an efficient targeting strategy to bypass the BBB and a judiciously designed nanocarrier with the desirable characteristics described above. The glycemic control strategy has only been reported for the delivery of small nucleic acids (siRNA or ASO) ^[7b, 13]^.

**Figure 1.**
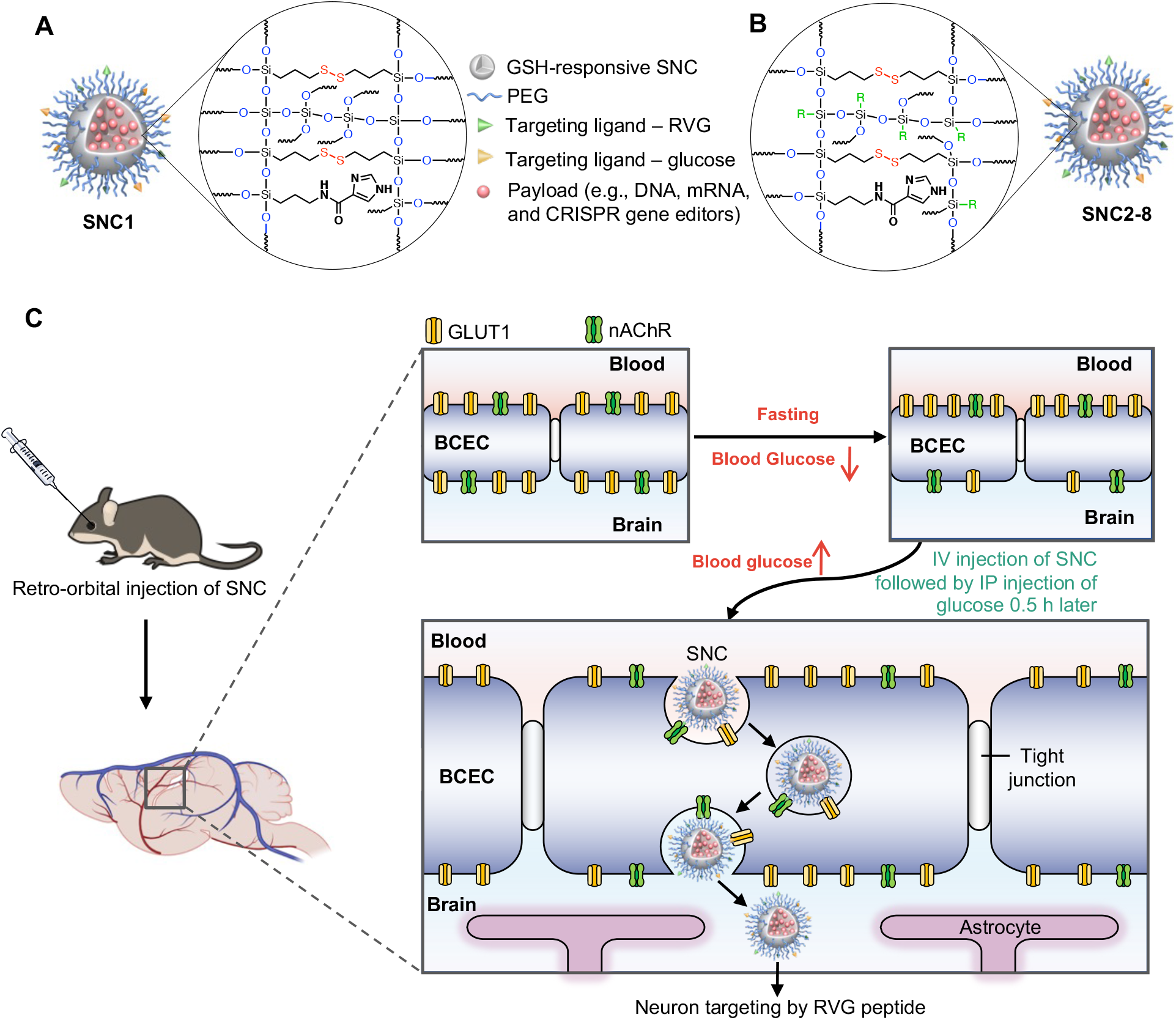
Design of the SNC formulation and brain targeting strategy. **(A-B)** Schematic illustration of SNC formed by silica precursor with (A) 4 active arms (i.e., SNC1), or (B) 3 active arms (i.e., SNC2-8). R= nonhydrolyzable inactive arm. SNC formulations are summarized in Fig. 2A. **(C)** Schematic illustration of systemic delivery of SNCs into the brain via the dual targeting ligand strategy.

For this study, we designed and evaluated a library of glutathione (GSH)-responsive silica nanocapsules (SNCs) (Fig. 1A) using different silica reagents (Fig. 2A) aimed at finding the optimal SNC formulation to achieve high delivery efficiencies for various biologics in the brain. The GSH-responsive SNCs were prepared using a water-in-oil microemulsion that allows for encapsulation of various types of biologics (i.e., DNA, mRNA, Cas9 RNP) with superior loading content and efficiency ^[14]^. The SNC is versatile both in terms of the type of payloads it can deliver and the type of surface ligands it can be conjugated with. These properties, in addition to its modular composition, make it a highly desirable nanoplatform for many applications including brain-targeted gene therapy and therapeutic genome editing. We demonstrated that intravenously injected SNCs with optimal composition and conjugated with glucose + RVG dual targeting ligands can bypass the BBB and efficiently deliver mRNA, Cas9 mRNA/sgRNA and Cas9 RNP into the brain, enabling efficient gene transfection and gene editing in the CNS. This is the first study on the brain-wide delivery of mRNA, Cas9 mRNA/sgRNA and Cas9 RNP via intravenous administration in healthy animals. We further investigated the potential of using brain-targeting SNC to edit two endogenous genes (i.e., amyloid precursor protein (*App*) and tyrosine hydroxylase (*Th*)) by the delivery of Cas9 mRNA and sgRNA. This brain-wide gene editing potential of SNC in wild-type mice broadens the potential application of gene therapy and therapeutic genome editing in treating various neurological diseases.

**Figure 2.**
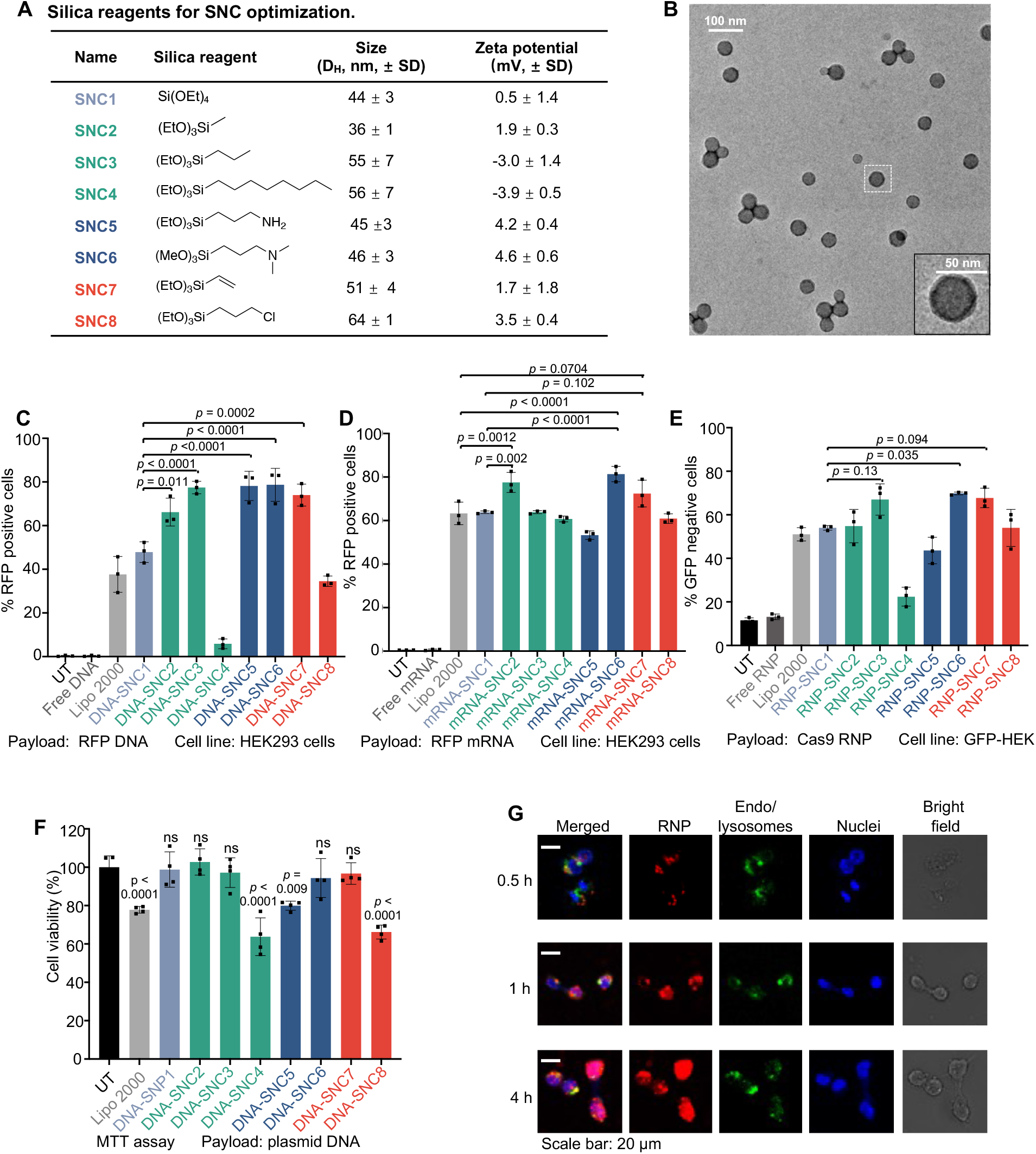
Optimization of the SNC formulation*in vitro*. **(A)** Summary of SNCs studied and the corresponding silica reagents. SNCs were formed by silica precursors with 4 active arms (i.e., SNC1) or 3 active arms (i.e., SNC2-8). The sizes and zeta-potentials were characterized by dynamic light scattering (DLS) using DNA-encapsulated SNCs (i.e., DNA-SNCs). **(B)** Representative TEM image of RNP-encapsulated SNC7 with surface PEGylation. **(C-D)** Transfection efficiency of the **(C)** DNA- and **(D)** mRNA-encapsulated SNCs in HEK293 cells (n=3). **(E)** Gene editing efficiency of RNP-loaded SNCs in GFP-expressing HEK 293 cells (n=3). Control groups include untreated (UT), free payload (e.g., free DNA in (C), free mRNA in (D), and free RNP in (E)), and Lipofectamine (Lipo) 2000 loaded with the corresponding payload. **(F)** Viability of HEK 293 cells treated with DNA-loaded SNCs and DNA-complexed Lipo 2000 (n=4). For the viability study, statistical difference was calculated between each group and untransfected cells (UT). **(G)**. Intracellular trafficking of SNC7 loaded with Atto550-tagged RNP. The intracellular localization of the Atto550-tagged RNP was studied at 0.5 h, 1 h, and 4 h post-treatment. Data are presented as mean ± SD. Statistical significance (*p*-value) was calculated via one-way ANOVA with a Tukey post hoc test.

## 2. Results

### 2.1 Design, fabrication and evaluation of a library of SNCs for efficient delivery of DNA, mRNA, and Cas9 RNP*in vitro*

SNCs were prepared by a water-in-oil microemulsion method ^[14]^. Different biologics (i.e., nucleic acids, genome editors) can be encapsulated inside the SNCs with a high loading content (i.e., around 10 wt%) and loading efficiency (> 90 %). An imidazole-containing silica reagent, *N*-(3-(triethoxysilyl)propyl)-1*H*-imidazole-4-carboxamide (TESPIC), was incorporated into the SNC to enhance its endosomal escape capability ^[15]^. A disulfide bond-containing crosslinker, bis[3-(triethoxysilyl)propyl]-disulfide (BTPD), was incorporated into the silica network to yield GSH-responsive SNC that can quickly release the cargo in the cytosol, where the GSH concentration is 2-10 mM. SNC1 was fabricated with tetraethyl orthosilicate (TEOS). TEOS has four ethoxy groups (-OC_2_H_5_) that can hydrolyze and form siloxane bonds (-Si-O-Si-) to yield a compact and stiff SNC structure (Fig. 1A). Replacing TEOS used for SNC1 with various types of silica precursors that have 3 hydrolyzable groups (i.e., active arms) and 1 nonhydrolyzable group (i.e., inactive arm, as shown in Fig. 1B) may reduce the crosslinking density and hence the stiffness of the resulting SNCs, which may affect the pharmacokinetics of the SNCs and their internalization by target cells ^[16]^. Moreover, inactive arms with different functional groups (e.g., saturated/unsaturated fatty chains of different lengths, or with charged moieties) may further alter their physical/chemical properties and affect their behaviors *in vitro* and *in vivo*. We built a library of SNCs (i.e., SNC2-8, as shown in Fig. 2A) based on this hypothesis. PEG-silane with glucose (Glu) or RVG targeting ligands (i.e., silane-PEG-Glu and silane-PEG-RVG) were synthesized (Fig. S1A and S1B, NMR spectra of the products are shown in Fig. S2-S4) and conjugated onto the SNC surface. SNCs formed by different silica reagents had similar hydrodynamic diameters and nearly neutral surface charges (Fig. S5 B-C), indicating that the components of SNC did not affect their sizes and levels of PEGylation notably. As shown in Fig. 2B, RNP-loaded SNC7 exhibited a spherical structure with an average size of 40 nm under Transmission electron microscopy (TEM)

The delivery efficiencies of different payloads (i.e., DNA, mRNA and RNP) by SNCs composed of silica reagents containing an inactive arm carrying different moieties (i.e., SNC2~SNC8) were compared with SNC1. SNCs exhibited distinct delivery performance for different payload types. For DNA delivery (Fig. 2C), a number of SNCs (i.e., DNA-SNCs) showed a higher transfection efficiency (up to two-fold) than the commercially available agent, Lipofectamine 2000 (Lipo 2000), including SNCs with short fatty chain inactive arms (i.e., DNA-SNC2 and DNA-SNC3), SNCs with charged inactive arms (i.e., DNA-SNC5 and DNA-SNC6), and SNC with an unsaturated inactive arm (i.e., DNA-SNC7). For mRNA delivery (Fig. 2D), mRNA-SNC2, mRNA-SNC6 and mRNA-SNC7 showed an increased transfection efficiency in comparison with Lipo 2000 and mRNA-SNC1. We then investigated the genome-editing efficiency of RNP-encapsulated SNCs (i.e., RNP-SNC) by delivering the RNP targeting the GFP gene in a transgenic GFP-expressing HEK 293 cell line. As shown in Fig. 2E, three SNC formulations, namely, the two SNCs with short fatty chain inactive arms (i.e., RNP-SNC2 and RNP-SNC3), and the SNC with tertiary amine inactive arm (i.e., RNP-SNC6), showed up to 1.6-fold higher gene editing efficiency than Lipo 2000 or RNP-SNC1. The MTT assay was performed to study the biocompatibility of DNA-SNCs (Fig. 2F). DNA-SNC1 and most of the SNC formulations (i.e., DNA-SNC2, DNA-SNC3, DNA-SNC6 and DNA-SNC7) with enhanced nucleic acid/genome editor delivery efficiencies exhibited neglectable cytotoxicity.

The storage stability of SNC was studied by dispersing DNA-encapsulated SNC7 in a storage buffer (i.e., 20 mM HEPES-NaOH pH 7.5, 150 mM NaCl, 10% glycerol) and stored at −80 oC. The transfection efficiency of SNC was tested every 7 days in HEK cells. As shown in Fig. S6A, SNC7 was stable for 17 weeks without significant transfection efficiency change. Surface PEGylation is a useful strategy to improve the colloidal stability of nanoparticles ^[17]^. The colloidal stability of SNC7 with or without targeting ligands (i.e., glucose and RVG) was studied in an aqueous solution (Fig. S6B). The hydrodynamic diameter of SNC7 remained unchanged for at least 60 h, indicating excellent colloidal stability of SNC due to the surface PEGylation.

The intracellular trafficking of SNC7 loaded with Atto550-tagged RNP was studied in HEK293 cells (Fig. 2G). RNP was mostly co-localized with endo/lysosomes (labeled by Lysotracker Green DND-26) 0.5 h after incubation. The RNP exhibited significantly higher level of co-localization with the nuclei (labeled by Hoechst 33342) and lower level of co-localization with endo/lysosomes 4 h post-treatment, indicating successful endosomal escape and nuclear transportation of RNP. To study the GSH-responsive behavior of SNC, SNC7 loaded with GFP mRNA was incubated with HEK293 and NIH 3T3 cells in culture media containing intentionally added GSH with a GSH concentration ranging from 0 to 10 mM. As shown in Figure S7, the mRNA transfection efficiency was not affected at GSH concentrations lower than 0.5 mM, suggesting that the SNC is stable in the extracellular space. However, a significant decrease in the mRNA transfection efficiency was observed at a GSH concentration of 0.5 mM or higher. The cytosol GSH concentration ranges from 1 to 10 mM, while the plasma/extracellular GSH concentration ranges from 1 to 20 μM ^[18]^. These results suggest that the SNC remains stable before internalization by cells, but it can effectively break down in the cytosol to readily release the payload.

### 2.2 Optimization of SNC formulation for brain-targeted delivery

We next investigated the *in vivo* RNP and mRNA delivery efficiency of SNCs in Ai14 mice. The Ai14 mouse genome contains a CAGGS promoter and a LoxP-flanked stop cassette (i.e., three SV40 polyA sequences), which prevents the expression of the downstream tdTomato gene. Cre recombinase-encoding mRNA (i.e., Cre mRNA), or Cas9 RNP targeting the SV40 polyA sequence can remove the stop cassette and lead to tdTomato expression in the targeted tissue/cells, which can be used to detect the transfection/gene editing efficiency *in vivo* (Fig. 3A and 3B) ^[5a, 14]^. However, the editing efficiency of RNP is significantly under-reported based on the tdTomato expression level using the Ai14 report mice, because tdTomato expression requires multiple RNP-mediated DNA breaks to generate excisions of at least two of three SV40 polyA sequences **(Fig. 3A and 3B)**^[5a]^.

**Figure 3.**
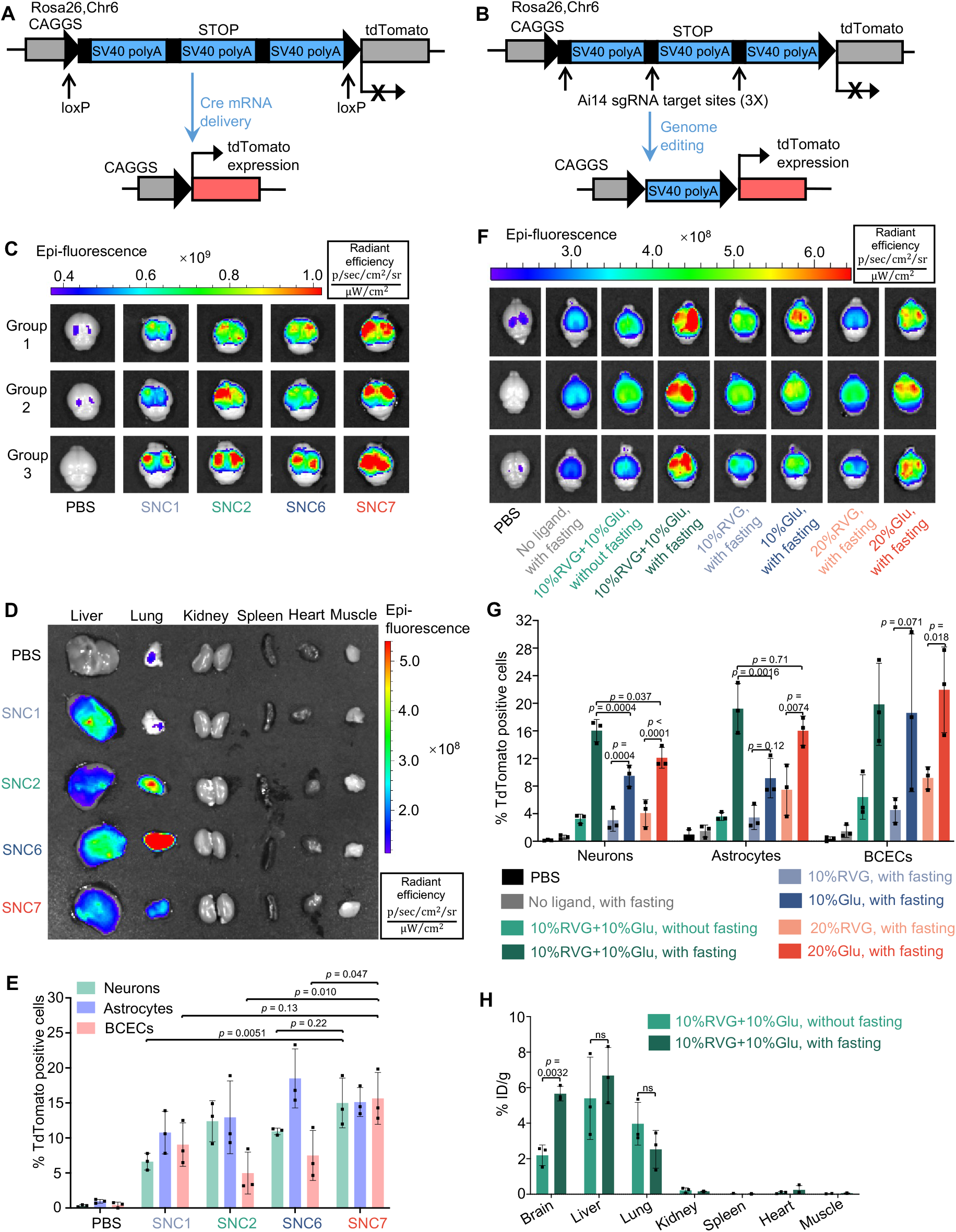
Optimization of SNC formulation and targeting ligand for brain delivery via systemic administration. **(A-B)** The tdTomato locus in the Ai14 reporter mouse. **(A)** TdTomato expression can be achieved by Cre mRNA through Cre-Lox recombination. **(B)** The stop cassette containing three Ai14 sgRNA target sites prevents downstream tdTomato expression. Excision of two SV40 polyA blocks by Ai14 RNP results in tdTomato expression. **(C-E)** Optimization of SNC formulation for brain delivery using Cre mRNA as a payload and 10%Glu+10%RVG as the brain targeting ligands (Cre mRNA dosage 2 mg/kg, n=3). **(C)** and **(D)**, *ex vivo* IVIS images of brains **(C)** and major organs **(D)** of Ai14 mice retro-orbitally injected with SNC1, SNC2, SNC6 and SNC7. **(E)** IFCM analysis of the editing efficiencies of three types of brain cells, namely, neurons, astrocytes, and BCECs (n=3). **(F-H)** Optimization of the targeting ligands for efficient brain delivery (Cre mRNA dosage 4 mg/kg, n=3). Cre mRNA-loaded SNC7 conjugated with the various amount (0 to 20%) and types (RVG or Glu alone or RVG+Glu hybrid) were IV injected into Ai14 mice with or without fasting. **(F)***Ex vivo* IVIS images of Ai14 mouse brains retro-orbitally injected with SNC7 conjugated with different types/amounts of targeting ligands and with or without glycemic control; **(G)** The effects of various brain targeting strategies on the in vivo editing efficiency of SNC7 in three types of brain cells analyzed by IFCM (n=3). **(H)** The effect of glycemic control on the biodistribution of SNC7-10%Glu+10%RVG loaded with Atto550-tagged RNP in wild-type mice (n=3). Data are presented as mean ± SD. Statistical significance (*p*-value) was calculated via one-way ANOVA with a Tukey post hoc test.

To study the effects of the SNC formulations on the delivery efficiency of mRNA *in vivo*, SNCs with higher *in vitro* mRNA delivery efficiencies than Lipo 2000 (i.e., SNC2, SNC6 and SNC7), as well as SNC1 were selected for *in vivo* evaluations. These four types of SNCs were conjugated with glucose and RVG with a feed molar ratio of mPEG-silane : Glu-PEG-silane : RVG-PEG-silane = 8 : 1 : 1 (i.e., 10%Glu+10%RVG). The Cre mRNA encapsulated SNCs were intravenously injected (i.e., retro-orbital injection; mRNA dose: 2 mg/kg) to Ai14 mice with glycemic control (i.e., 24 h fasting prior to SNC injection and intraperitoneal injection of glucose 0.5 h after SNC injection). While all four Cre mRNA-encapsulated SNC (i.e., mRNA-SNC) formulations showed tdTomato expression in the brain, SNC7 (i.e., with unsaturated inactive arm) exhibited the highest tdTomato fluorescence intensity (Fig. 3C). All the SNCs showed moderate Cre mRNA delivery in the liver, while SNC2, SNC6 and SNC7 also showed tdTomato expression in the lungs (Fig. 3D and Fig. S8A). Interestingly, although SNC6 showed the highest mRNA delivery efficiency *in vitro*, it only exhibited moderate mRNA delivery efficiency in the brain; instead, it had high accumulation in the lungs and induced strong tdTomato expression in pulmonary endothelial cells and maybe alveolar epithelial cells per the immunofluorescence staining (Fig. 3D and Fig. S9), demonstrating the different behaviors of SNCs in blood circulation. The superior mRNA delivery efficiency of SNC7 in the brain was confirmed by immunofluorescence staining. Brains were fixed, cryosectioned and stained with anti-tdTomato antibody to confirm tdTomato expression. The coronal section mosaic tile image of the mouse brain showed tdTomato expression in the whole brain (Fig. S8B). Notably, SNC7 induced wide-spread and high-level tdTomato expression, especially in the cortex and hippocampus, suggesting the spread of SNCs in the brain parenchyma after bypassing the BBB. The editing efficiencies of three types of brain cells were evaluated by immunofluorescence flow cytometry (IFCM), using primary antibodies against cell type-specific markers (i.e., NeuN for neurons, GFAP for astrocytes, and CD31 for BCECs) and the corresponding fluorophore-tagged secondary antibodies (Fig. 3E, gating strategies of IFCM are shown in Fig. S10-S12). SNC7 exhibited the highest editing efficiency in the neuron (15%) and was selected for further studies.

### 2.3 Optimization of the targeting ligands for brain-targeted delivery

We hypothesized that dual targeting ligands (i.e., the combination of Glu + glycemic control and RVG) can further enhance the capability of the SNCs bypassing the BBB (Fig. 1C). To prove this hypothesis, Cre mRNA-SNC7 conjugated with various amounts (0, 10, or 20%) and types (RVG alone, glucose alone, or a combination of RVG and glucose) of surface ligands were intravenously injected into Ai14 mice with or without glycemic control (mRNA dosage: 4 mg/kg). Among all the groups studied, Cre mRNA-loaded SNC7-10%RVG+10%Glu with glycemic control exhibited the highest tdTomato expression in the brain (Fig. 3F). Meanwhile, its accumulation and mRNA transfection efficiency in other organs (e.g., the lungs and liver) were relatively low (Fig. S13). The mRNA transfection/gene-editing efficiencies in specific brain cell types were quantified by IFCM (Fig. 3G). Consistent with the IVIS data, SNC7-10%RVG+10%Glu exhibited significantly higher neuron (16%) and astrocyte (19%) editing efficiencies than single-ligand formulations. Without fasting, SNC-10%RVG+10%Glu showed a similar brain delivery efficiency to SNCs with only RVG conjugation, indicating that glycemic control can efficiently boost the BBB-crossing capability of glucose-conjugated SNC. The biodistribution of SNC7-10%RVG+10%Glu was quantified using Cas9 RNP with an Atto550-tagged sgRNA (RNP dose: 5 mg/kg). The Atto550 fluorescence intensity in major organs was quantified 24 h post-injection. As shown in Fig. 3H, SNC7-10%RVG +10%Glu-treated mice with glycemic control exhibited a 2.6-fold higher brain accumulation than the ones without glycemic control (i.e., 5.7% vs. 2.2 % ID/g). Meanwhile, no significant difference was observed in other organs such as the liver (6.6% vs. 5.4% ID/g) and the lungs (2.5% vs. 4.0% ID/g) The best-performing formulation, SNC7 with 10%Glu+10%RVG targeting ligand combination, will be referred to as SNC from here on.

### 2.4 Efficient brain-wide delivery of mRNA and RNP by SNC

With the optimized SNC formulation and targeting strategy, we then evaluated the brain-targeting delivery efficiency of Cas9 RNP in Ai14 reporter mice (Cas9 RNP dosage: 5 mg/kg). The effect of multiple injections (i.e., triple (3×) injection versus single (1×) injection) was also investigated using Cre mRNA in Ai14 mice (Cre mRNA dosage for each injection, 4 mg/kg). The timeline of triple injection is shown in Fig. 5F. Coronal section mosaic tile image of representative brain slices (Fig. 4A-E,+1.18 mm bregma; Fig. 4F-J, −1.70 mm bregma) clearly showed tdTomato expression in the whole brain by systemic delivery of RNP and Cre mRNA. Furthermore, triple injection of Cre mRNA-encapsulated SNC (Fig. 4E and 4J) induced much higher tdTomato expression than single injection (Fig. 4D and 4I).

**Figure 4.**
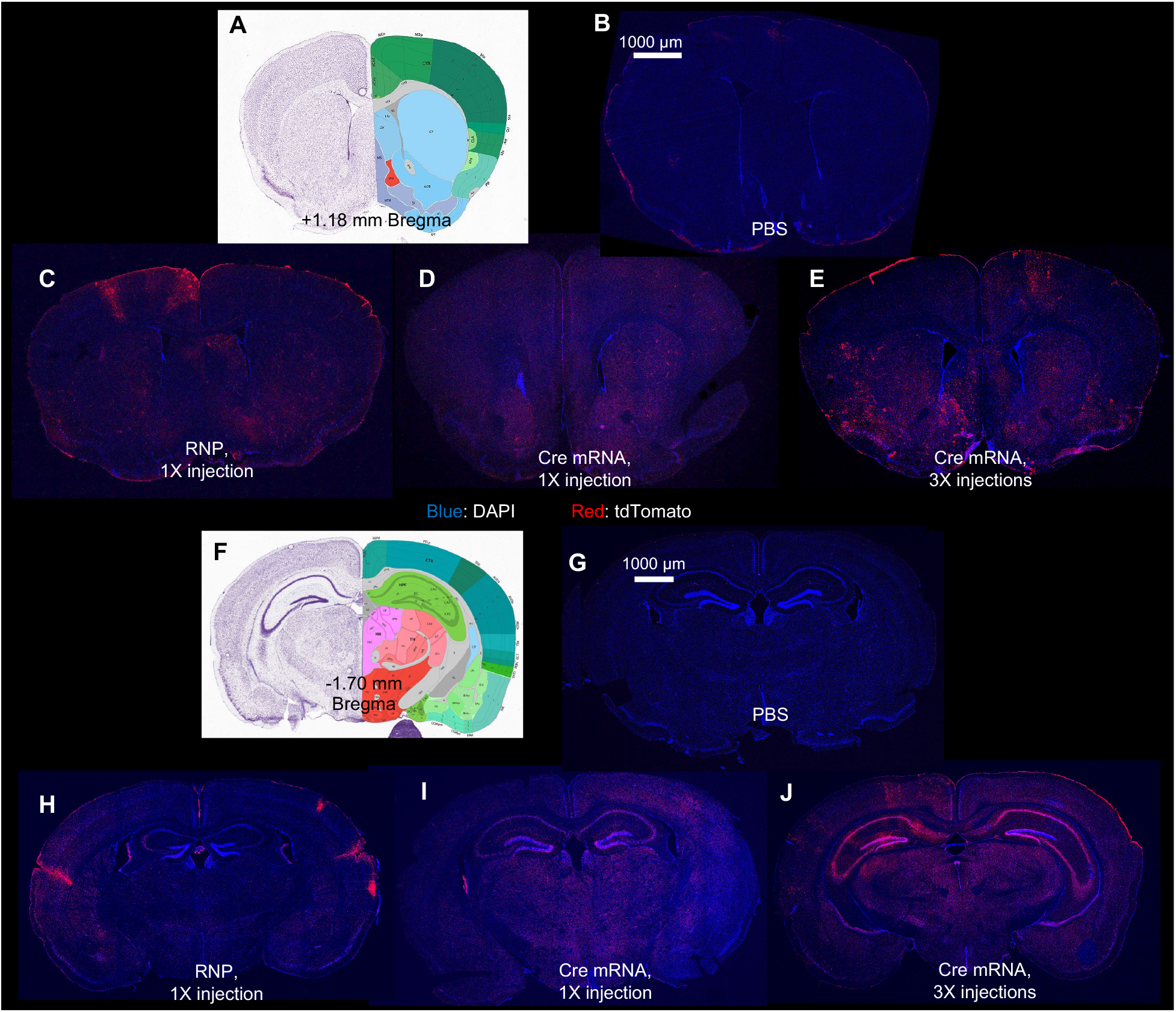
RNP- and Cre mRNA-encapsulated SNC7-10%Glu+10%RVG achieved whole-brain editing. Representative coronal section mosaic tile CLSM images of the brains of Ai14 mice intravenously injected with PBS, RNP-encapsulated SNC (1× injection), and Cre mRNA-encapsulated SNCs (1× and 3× injections) at +1.18 mm bregma **(B-E)** and −1.70 mm bregma **(G-J)**. Coronal cartoons of the mouse brain were cited from ^[29]^. Blue, DAPI staining nuclei; red, tdTomato.

**Figure 5.**
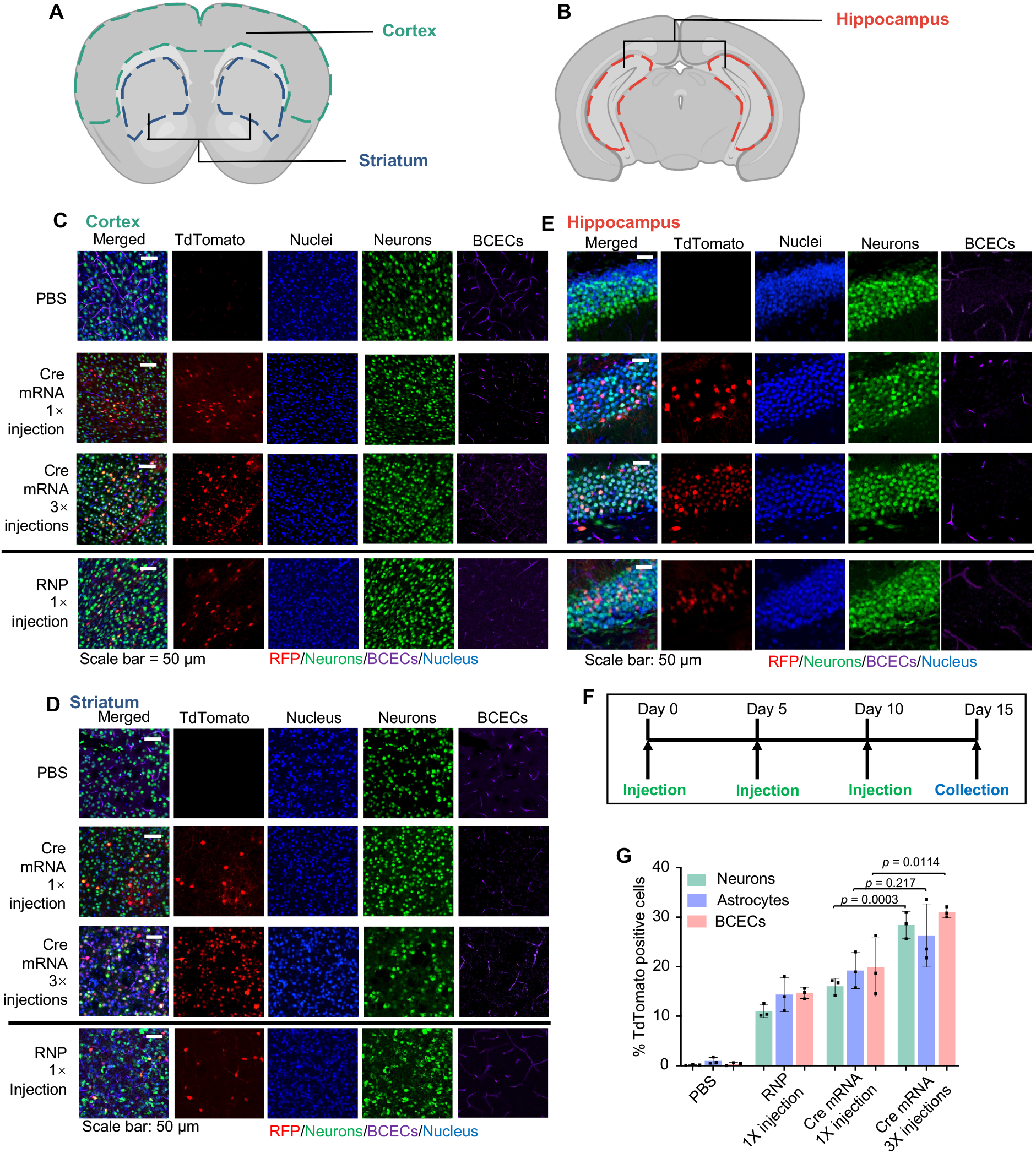
Cre mRNA- and Cas9 RNP-encapsulated SNCs induced tdTomato expression in neurons and other brain cell types. **(A-B)** Regions of interest for CLSM imaging (coronal view), colored dash lines enclose the cortex, striatum **(A)** and hippocampus **(B)**. **(C-E)** Representative CLSM images of **(C)** cortex, **(D)** striatum, and **(E)** hippocampus with tdTomato positive cells co-localizing with cell type-specific markers. Scale bar: 50 μm. **(F)** Timeline for SNC triple (3×) injections. **(G)** IFCM analysis of tdTomato positive neurons, astrocytes, and BCECs. Data are presented as mean ± SD (n=3). Statistical significance (*p*-value) was calculated via one-way ANOVA with a Tukey post hoc test.

We next performed triple-color immunofluorescence staining and confocal laser scanning microscopy (CLSM) to visualize the cell types of interest with tdTomato expression in different brain regions including cortex, striatum, and hippocampus (Fig. 5A and 5B). TdTomato expression was confirmed in all brain regions. The majority of tdTomato-positive cells were overlapping with NeuN positive cells (neurons), while a small portion of the tdTomato-positive cells were overlapping with CD31 positive cells (BCECs) (Fig. 5C-E). The immunofluorescence staining of astrocytes using anti-GFAP antibody also revealed edited astrocytes in different brain regions (Fig. S14). The transfection/gene-editing efficiencies in three types of brain cells were quantified by IFCM. For single injection of RNP-encapsulated SNC, about 11% of neurons, 14% of astrocytes, and 15% of BCECs were edited. For single- and triple-injection of Cre mRNA encapsulated SNC, the corresponding transfection/editing efficiency for neurons/astrocytes/BCECs were 16%/19%/20% and 28%/26%/31%, respectively (Fig. 5G), further confirming that higher transfection/editing efficiency can be achieved via multiple injections.

### 2.5 Brain-wide delivery of SNC induces editing of disease-relevant genes

Many neurodegenerative diseases (NDDs) including Alzheimer’s disease (AD) and Parkinson’s disease (PD) are linked with gene mutations ^[2c, 19]^. The CRISPR/Cas9 system can target and edit disease-causing mutations in a sequence-dependent manner, resulting in a permanent genetic change and thus offering the promise to treat the root causes of genetic diseases ^[1]^. To assess the gene editing efficiency of SNCs, we chose two endogenous gene targets, i.e., the *App* gene related to AD, and the *Th* gene related to PD.

For *App* gene editing, we used a recently reported, unique *App* sgRNA (aka *App*^659^ sgRNA) targeting the C-terminus of APP encoding the cleavage site for β-secretase 1 (i.e., BACE-1) and reciprocally manipulating the amyloid pathway ^[20]^. This *App* targeting strategy works by limiting APP and BACE-1 approximation ^[20-21]^, which can attenuate APP-β-cleavage and Aβ production while up-regulating neuroprotective APP-α-cleavage (Fig. S15) ^[2c, 20]^. Using this *App* targeting strategy, the APP N-terminus and compensatory APP-homologs remain intact. This selective APP silencing strategy enabled by SNC may lead to safe and non-invasive gene therapy for AD. We demonstrated that SNC loaded with Cas9 mRNA and *App*^659^ sgRNA (weight ratio of Cas9 mRNA: sgRNA at 4:1) can efficiently edit *App* in NIH 3T3 fibroblasts (Fig. S16). SNC loaded with Cas9 mRNA and *App*^659^ sgRNA (i.e., *App*-SNC) was intravenously injected into C57BL/6J wild-type mice following the triple-injection timeline (Fig. 5F). The total RNA dosage for this study was 5 mg/kg per injection. The gene editing efficiencies of *App*-SNC in different regions of the cerebrum were quantified by next-generation sequencing (NGS). The *App* gene editing efficiency for the cortex, hippocampus, and thalamus/hypothalamus was 3.6%, 3.4 % and 6.1 %, respectively (Fig. 6A-B, Fig. S17A-C). *App* gene editing also occurred in the liver (6.8%) and lungs (2.7%). The editing spectra (Fig. S17C) contained a 5-base deletion as the most frequently mutated *App* loci resulting from *App*^659^ sgRNA-induced gene editing, consistent with a previous report using this sgRNA ^[22]^. The disruption of the APP C-terminus was also confirmed at the protein level using a C-terminus-specific antibody Y188. The reduction of intact APP in the cerebrum was 19.1 % as analyzed by western blot (Fig. 6C-D). Immunofluorescence staining of brain slices also showed an obvious reduction of intact APP that can be recognized by Y188 antibody, in different brain regions including the hippocampus and cortex (Fig. 6E-F, CLSM images of the thalamus in Fig. S18), indicating *App*-SNC can effectively modulate APP *in vivo*. Brain-wide gene editing was also observed using SNC targeting the *Th* gene (i.e., *Th*-SNC). The editing efficiency of *Th*-SNC, quantified by NGS, in the cortex, hippocampus and thalamus/hypothalamus was 3.9%, 3.4%, and 3.5%, respectively (Fig 6G, Fig. S18D-F). *Th* gene editing was also observed in other major organs (e.g., 3.0% in the liver and 3.3% in the lungs). Impressively, a significant reduction in the TH protein level of the cerebrum (i.e., 30.3%), analyzed by western blot, was observed (Fig. 6H-I). Immunofluorescence staining of brain slices also showed significant TH reduction in major TH-expression areas including the ventral tegmental area and substantia nigra (Fig. 6J-L, coronal section mosaic tile CLSM images in Fig S19). These results confirmed SNC with the optimal formulation can efficiently deliver gene editors and produce gene editing in the whole brain.

**Figure 6.**
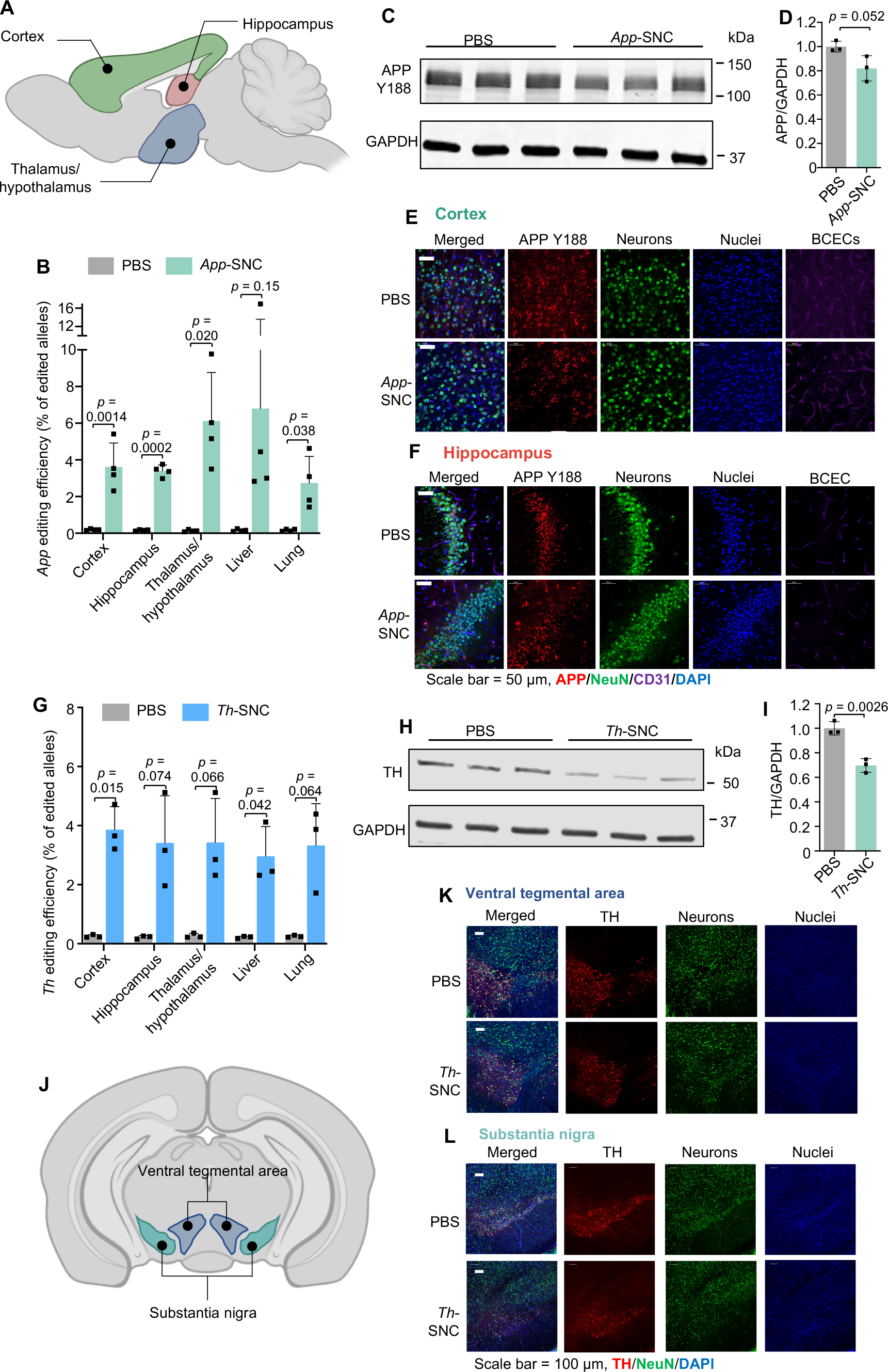
Systemic delivery of Cas9 mNRA+sgRNA encapsulated SNCs induced disease-relevant endogenous genes in wild-type mouse brain. **(A-F)** APP C-terminus editing by *App*-SNC via intravenous injection. **(A)** An illustration of brain regions collected for NGS (sagittal view). **(B)***App* editing efficiency in different brain regions and major organs by NGS after intravenous injection (n=4). **(C-F)** Brains of *App*-SNC treated mice were immunostained by APP Y188 antibody recognizing the APP C-terminus and analyzed by western blot (n=3) (**C-D**) and CLSM (**E-F**). **(G-L)***Th* gene editing by *Th*-SNC in wild-type mice via intravenous injection. **(G)***Th* editing efficiency in different brain parts and major organs by NGS (n=3). **(H-I)** Brains of *Th*-SNC treated mice were immunostained by an anti-TH antibody and analyzed by western blot (n=3). **(J)** An illustration of brain regions of interest (i.e., TH-expressing regions) for CLSM imaging (coronal view). (K-L) Representative CLSM images of **(K)** ventral tegmental area, and **(L)** substantia nigra with TH and NeuN immunofluorescence staining. Scale bar: 100 μm. Data are presented as mean ± SD. Statistical significance (*p*-value) was calculated via one-way ANOVA with a Tukey post hoc test.

### 2.6 Biocompatibility evaluation of SNC

The systemic toxicity of *App*-SNC was first investigated by blood biochemical analysis (Fig. S20). Serum was collected from wild-type mice that received PBS or *App*-SNC triple injections and a panel of blood biochemical parameters was measured. No significant variations were found in these biochemical parameters. The systemic toxicity was also evaluated by hematoxylin and eosin (H&E) staining. H&E staining was performed using the tissue slices including the liver, lungs, spleen, kidneys, and heart (Fig. S21). There was no evidence of histological change or inflammatory cell infiltration in these tissues, indicating good biocompatibility of SNC.

To study the toxicity and immunogenicity of SNC in the target organ (i.e., the brain), brain slices of 3× *App*-SNC injected wild-type mice were H&E stained and compared with PBS control mice (Fig. S22). Again, no sign of toxicity or immunogenicity was found associated with SNC injection in different brain regions (i.e., cortex, striatum, and hippocampus). To analyze the immune response, mRNA from brain cells was extracted and the relative expression levels of a panel of inflammatory cytokines were evaluated by reverse transcription quantitative PCR (RT-qPCR). As shown in Fig. S23, except for a slight elevation of TNF-α, the expression levels of all other cytokines analyzed remain unchanged in *App*-SNC-injected mice, suggesting good biocompatibility.

## 3. Discussion

The promise of gene therapy in the brain relies on the safe and efficient delivery of biologics (e.g., DNA, mRNA, and CRISPR genome editors) to the brain, which is extremely challenging due to the blood-brain barrier as well as the dense brain tissue ^[10c, 23]^. In this work, we have demonstrated that SNC conjugated with glucose and RVG dual targeting ligands can effectively bypass intact (not leaky) BBB and efficiently deliver diverse biologics including CRISPR genome editors and nucleic acids to the whole brain via systemic administration. Previously reported non-viral vectors, designed for the delivery of biologics to the brain, often rely on leaky vasculature or compromised BBB caused by certain diseases such as stroke and glioblastoma ^[24]^. Innovative strategies enabling the nanocarriers to (1) effectively bypass functional and intact BBB, (2) subsequently diffuse into the brain parenchyma, and (3) efficiently deliver the biologics to the brain cells are urgently needed for the prevention and treatment of various neurological genetic diseases in their early stages before BBB is compromised. The family of versatile SNCs we engineered offer versatile surface chemistry to enable the conjugation of various types of targeting ligands including the brain/neuron-targeting ligands used in this study. Varying the types of silica precursors used to prepare the SNCs yields SNCs with different *in vitro* (Fig. 2H-G)and *in vivo* delivery efficiencies and biocompatibility (Fig. 3C-E, Fig. S17-19). The versatility offered by this unique SNC nanoplatform in terms of its composition, morphology, surface chemistry, and types of payloads makes it highly desirable for many potential applications targeting different organs (via surface targeting ligand conjugation) and different diseases (via a judicious selection of the payload). We have discovered that a dual brain-targeting strategy (i.e., glucose and RVG targeting ligands) can efficiently enable the NPs to cross the BBB and distribute throughout the whole brain (Fig. 3-5). Also, for the first time, we demonstrated that mRNA, Cas9 mRNA/sgRNA, and Cas9/sgRNA RNP can be efficiently delivered by SNCs to the whole brain of healthy mice with intact BBB systemically via a non-viral approach. SNCs loaded with genome editors (i.e., Cas9 mRNA/sgRNA targeting *App* or *Th* gene) were able to edit disease-relevant endogenous genes brain-wide in healthy wild-type mice, leading to 19.1% reduction in the expression level of intact APP and 30.3% reduction in the expression level of TH (Fig. 6). A previous study has shown that about 20% decrease in the protein level (e.g., BACE1 knockdown via siRNA therapy, 7 injections) in AD model mice can lead to phenotype/behavioral changes ^[7a]^. Another study has shown that systemic delivery of anti-BACE1 siRNA in wildtype mice (1mg/kg, two injections) resulted in a 20% reduction in protein; while repeated injections in AD model mice (1 mg/kg, 10 injections) induced a more significant reduction in protein level compared to the PBS control or scrambled siRNA control groups (30% in the hippocampus, and 50% in the cortex), leading to significant phenotype and behavioral improvements^[13]^. Considering siRNA-mediated therapy is transient, permanent gene editing enabled by CRISPR genome editors delivered by our unique SNC nanoplatform, boosted by multiple injections and compromised BBB in AD animals/patients ^[8c, 25]^, will produce a more significant change in the protein level and likely lead to significant behavioral and phenotype improvement.. Thus, our results demonstrate that SNC as a versatile non-viral delivery nanoplatform for genome editors has a great potential for treating neurological diseases safely and non-invasively.

One limitation of the current approach is the off-target delivery of biologics to other organs (e.g., liver and lung) or other brain cell types (e.g., astrocytes and BCECs), instead of specific delivery to the neurons in the brain. This might not be a concern if the therapeutic gene editing is aimed at disrupting the inherited gene mutations as such genetic mutations occur in the whole body. For certain types of neurodegenerative diseases, gene editing in peripheral organs in additional to the central nervous system might even be beneficial. For example, the amyloid precursor protein (APP) we edited in this work, is known to be expressed in other types of brain cells (e.g., astrocytes and microglia) and peripheral tissues (e.g., liver, pancreas, adipose tissue, and myotubes) ^[26]^. Therapeutic gene editing of APP in both central nervous system and peripheral organs may be an effective strategy for the prevention and treatment of Alzheimer’s disease as well as other metabolic diseases ^[26a, 27]^. Additionally, to achieve cell specific genome editing, certain genome editors such as plasmid DNA encoding Cas9 protein and sgRNA with cell-specific promoters (such as neuron specific promoters) can be encapsulated into the SNC and achieve cell specific genome editing ^[28]^. As a proof-of-concept study, we have demonstrated that the glucose/RVG dual targeting strategy can significantly enhance the SNC’s ability to bypass intact BBB and deliver versatile biologics to the whole brain via systemic administration. Future studies will be focused on analyzing the effects of off-target delivery in other organs/cell types, as well as developing cell-type specific gene editing therapy for neurodegenerative diseases.

## 4. Conclusion

We have developed a family of biocompatible and versatile SNCs capable of delivering diverse biologics including CRISPR genome editors and nucleic acids. When the top-performing SNC (SNC7) was conjugated with 10% glucose and 10% RVG dual brain targeting ligands, it can efficiently bypass the BBB and generate high transfection and/or genome editing efficiency in the whole brain via systemic delivery. This unique SNC nanoplatform will lead to new, safe, efficient, and non-invasive genome editing therapies to treat the root causes of genetic CNS disorders. Given the modularity and versatility of the SNCs including their ability to encapsulate and deliver various types of CRISPR genome editors (e.g., base editors and prime editors), and the ease of targeting different genes by the CRISPR system, we anticipate that our SNCs will be applicable for the treatment of a wide range of neurological diseases, including neurodevelopmental disorders (e.g., Fragile X syndrome and Rett syndrome) and neurodegenerative diseases (e.g., Parkinson’s and Huntington’s diseases).

Future research will focus on (1) further optimization of the SNC including the amounts of glucose/RVG ligands conjugated onto the surface of SNC in order to further enhance their brain targeting abilities, and (2) testing the therapeutic efficacy and biosafety of the SNC nanosystem in neurological disease models (e.g., AD mouse models).

## Acknowledgments

The authors would like to acknowledge the financial support from the University of Wisconsin−Madison and the Wisconsin Institute for Discovery. Part of Fig. 1B, 5A, 5B, 6A and 6J was created using materials from Biorender.com. The brain atlas in Fig. 4A and 4F were cited from mouse.brain-map.org.

## Data availability

The data that support the findings of this study are available from the corresponding author upon reasonable request.

