## Supplementary material for "Overcoming the Blood-Brain Barrier for Gene Therapy via Systemic Administration of GSH-Responsive Silica Nanocapsules": Methods, Supplementary Figures S1-S23, Supplementary Tables S1-S4

### Materials

All of the silica reagents listed in Fig. 2A, unless otherwise specified, were purchased from Tokyo Chemical Industry Co., Ltd., USA. Tetraethyl orthosilicate (TEOS), 1*H*-imidazole-4-carboxylic acid, thionyl chloride ($SOCl_2$), Triton X-100, dichloromethane (DCM), trifluoroacetate (TFA), (1-ethyl-3-(3-dimethylaminopropyl)carbodiimide hydrochloride) (EDC), N,N-diisopropylethylamine (DIPEA), 1-hydroxybenzotriazole hydrate (HOBt), tris(2-carboxyethyl)phosphine hydrochloride (TCEP) and ammonia (30% in water) were purchased from Fisher Scientific, USA. Hexanol, cyclohexane, and (3-aminopropyl)triethoxysilane (APTES) were bought from Tokyo Chemical Industry Co., Ltd., USA. Triethylamine (TEA) and dimethyl sulfoxide (DMSO) were purchased from Alfa Aesar, USA. Diethyl ether and acetone were purchased from Sigma-Aldrich, USA. Bis[3-(triethoxysilyl)propyl]-disulfide (BTPD) was obtained from Gelest, Inc., USA. Methoxy-poly(ethylene glycol)-silane (mPEG-silane, $M_n$ = 5000) and maleimide-poly(ethylene glycol)-silane (Mal-PEG-silane, $M_n$ = 5000) were purchased from JenKem Technology, USA. Carboxylate-poly(ethylene glycol)-silane (HOOC-PEG-silane, $M_n$ = 5000) was purchased from Nanocs Inc., USA. 1,2-*O*-Isopropylidene-*α*-*D*-glucofuranoside was purchased from Santa Cruz Biotechnology, USA. RVG peptide with a C-terminal cysteine (sequence: YTIWMPENPRPGTPCDIFTNSRGKRASNGC) was purchased from GenScript Biotech Corporation, USA. Nuclear localization signal (NLS)-tagged *Streptococcus pyogenes* Cas9 nuclease (sNLS-*Sp*Cas9-sNLS, i.e., Cas9) was obtained from Aldevron, USA. The sgRNA protospacer sequences were summarized in Table S1.

**Synthesis of *N*-(3-(triethoxysilyl)propyl)-1*H*-imidazole-4-carboxamide (TESPIC)**

A mixture of 1*H*-imidazole-4-carboxylic acid (500 mg, 3.8 mmol) and $SOCl_2$ (10 mL) was refluxed at 75 $^{o}$C overnight. The reaction mixture was then cooled down to room temperature and added into anhydrous toluene (30 mL). The precipitate was collected by filtration and vacuum-dried to yield the chloride intermediate, 1*H*-imidazole-4-carbonyl chloride. The chloride intermediate product was then suspended in anhydrous THF (10 mL), followed by the addition of triethylamine (500 mg, 5 mmol) and APTES (850 mg, 4mmol). The mixture was stirred at room temperature overnight under a nitrogen atmosphere, and then filtered. The filtrate was collected and the solvent was removed by rotary evaporation to yield the final product TESPIC. Since silica reactants tend to undergo hydrolysis/polymerization during column purification, TESPIC was synthesized and used without purification [1]. $^{1}$H NMR (400 MHz, DMSO-*D6*): δ 0.65 (dd, 2 H), δ 1.15 (t, 9 H), δ 1.65 (dt, 2 H), δ 2.75 (m, 2 H), δ 3.85 (q, 6 H), δ 7.05 (s, 1 H), δ 7.43 (s, 1 H). $^{13}$C NMR (100 MHz, DMSO-*D6*): δ 167, 136, 135, 126, 60, 42, 22, 19, and 8 ppm.

**Synthesis of silane-PEG-Glucose (Glu)**

The synthesis scheme of glucose conjugated PEG-silane is shown in Fig. S1A, it is a three-step reaction.

**Synthesis of 3,5-*O*-benzylidene-1,2-*O*-isopropylidene-*α*-*D*-glucofuranoside (BIG, intermediate reactant 1)**. The intermediate reactant, 3,5-*O*-benzylidene-1,2-*O*-isopropylidene-*α*-*D*-glucofuranoside (**1**) was synthesized as reported previously [2]. Briefly, 3,5-*O*-benzylidene-1,2-*O*-isopropylidene-*α*-*D*-glucofuranoside (5 g, 23 mmol) and anhydrous zinc chloride (6.8 g, 50 mmol) were mixed in benzaldehyde (20 mL, 198 mmol) at room temperature under vigorous stirring overnight. The mixture was then dissolved in ethyl acetate (50 mL) and washed three times with DI water (50 mL). The organic phase was collected, and dried over anhydrous sodium sulfate. The organic solvent was subsequently removed by a rotary evaporator. The crude product was recrystallized from hexane to yield the final product as a white solid. $^1$H-NMR spectrum of the product (1) was shown in Fig. S2 ($CDCl_3$, 400 MHz).

**Synthesis of Silane-PEG-BIG (2)**. BIG was conjugated to silane-PEG-COOH via an esterification reaction. BIG (92.4 mg, 0.3 mmol) and silane-PEG-COOH (1 g, 0.2 mmol) were dissolved in anhydrous DMF (10 mL), followed by the addition of EDC (62 mg, 0.4 mmol), HOBt (54 mg, 0.4 mmol) and DIPEA (78 mg, 0.6 mmol). The mixture was stirred at room temperature overnight and precipitated with diethyl ether (100 mL) to obtain the intermediate product silane-PEG-BIG without further purification.

**Synthesis of Silane-PEG-Glu (3)**. Silane-PEG-BIG (150 mg) was dissolved in TFA/$CH_2Cl_2$ (1:1, v/v, 5 mL) and stirred for 1 h at room temperature. The solvent was then removed by a rotary evaporator. The residual was dissolved in anhydrous DMF (3 mL) and precipitated with diethyl ether (30 mL). The precipitate was washed twice with diethyl ether, and dried under vacuum to obtain the product silane-PEG-Glu without further purification. $^1$H-NMR spectrum of the product (3) is shown in Fig. S3 (DMSO-$D_6$, 400 MHz).

**Synthesis of Silane-PEG-RVG**

Silane-PEG-RVG was synthesized by a maleimide-thiol Michael addition reaction. RVG peptide (340 mg, 0.1 mmol) was mixed with silane-PEG-Mal (500 mg, 0.1 mmol) and dissolved in anhydrous DMF (10 mL). TCEP (57.4 mg, 0.2 mmol) was added to the mixture, and the reaction was carried out at room temperature for 16 h in a nitrogen atmosphere. The product, silane-PEG-RVG, was purified by precipitation with diethyl ether (50 mL) three times and dried under vacuum. The product was directly used for experiments without further purification. $^1$H-NMR spectrum of the product is shown in Fig. S4 (DMSO-$D_6$, 400 MHz).

**Preparation of the GSH-Responsive Silica Nanocapsules (SNCs)**

SNCs were synthesized by a water-in-oil microemulsion method as reported previously [3]. The oil phase was prepared by mixing Triton X-100 (1.77 mL) with hexanol (1.8 mL) and cyclohexane (7.5 mL). An aliquot of aqueous solution (30 μL) containing the desired payload (e.g., DNA, mRNA, or Cas9 RNP, 2 mg/mL) was mixed with the desired silica reagents (4 μL) (as shown in Table 1), BTPD (6 μL) and TESPIC (1 mg, 3 μmol). This mixture was homogenized by pipetting and then added to the oil phase (1.2 mL) The water-in-oil microemulsion was formed by vortexing for 1 min. Under vigorous stirring (1,500 rpm), an aliquot of 30% aqueous ammonia solution (4 μL) was added and the water-in-oil microemulsion was stirred at 4 °C for 12 h to obtain unmodified SNCs. Acetone (1.5 mL) was added to the microemulsion to precipitate the SNCs, the precipitate was recovered by centrifugation and was subsequently washed twice with ethanol and three times with water. The purified SNCs were finally collected by centrifugation.

The as-prepared, unmodified SNC was re-dispersed in DI water (3 mL). For surface modification, mPEG-silane, or a mixture mPEG-silane + silane-PEG-targeting ligands with different molar ratios were added to the above-mentioned SNC suspension. The total amount of PEG is 10 wt% of SNC. The pH of the suspension was adjusted to 8.0 using a 30% aqueous ammonia solution. The mixture was stirred at room temperature for 4 h. The resulting SNCs were purified by washing with DI water three times and were subsequently concentrated with Amicon Ultra Centrifugal Filters (Millipore Sigma, USA).

**Characterization**

The chemical structures of the products were confirmed by nuclear magnetic resonance (NMR) spectroscopy (Avance 400, Bruker Corporation, USA). The hydrodynamic diameters and zeta potentials of the SNCs were characterized by a dynamic light scattering (DLS) spectrometer (Malvern Zetasizer Nano ZS) at a 90° detection angle with a sample concentration of 0.1 mg/mL. SNCs were resuspended in DI water and the pH was adjusted to 7.4 for DLS analysis.

**Cell Culture for *In Vitro* Studies**

Human embryonic kidney cells (i.e., HEK 293 cells) were used for *in vitro* studies. HEK293 cells were obtained from ATCC (USA). GFP-expressing HEK 293 cells (i.e., GFP-HEK) were purchased from GenTarget Inc. NIH 3T3 fibroblast cells (i.e., 3T3 cells) were purchased from ATCC (USA). All cells were cultured in a medium containing DMEM (Gibco, USA) with 10% (v/v) fetal bovine serum (FBS, Gibco, USA) and 1% (v/v) penicillin-streptomycin (Gibco, USA) at 37 $^{o}$C with 5% carbon dioxide at 100% humidity.

**DNA and mRNA Transfection Efficiency Study**

An RFP-encoding plasmid DNA (i.e., RFP-DNA, Addgene #40260, USA) and an RFP-mRNA (Trilink Biotechnologies #L-7203, USA) were used for *in vitro* DNA and mRNA transfection studies, respectively. HEK 293 cells were seeded in 96-well plates 24 h prior to treatment, at a density of 10,000 cells/well. Cells were incubated with DNA-loaded SNCs or mRNA-loaded SNCs. A commercially available transfection agent, Lipofectamine 2000 (Lipo 2000) was used as the positive control. The dosage of DNA or mRNA was 200 ng/well. The Lipo 2000-DNA (or Lipo 2000-mRNA) complex was prepared following the protocols provided by the manufacturer, with a final dosage of Lipo 2000 at 0.5 μL per well. Untreated cells were used as the negative control. After 48 h, cells were harvested with 0.25% trypsin-EDTA, spun down and resuspended in 500 μL PBS. RFP expression efficiencies were obtained with a flow cytometer and analyzed with FlowJo 7.6. To study the stability of DNA encapsulated SNC in the presence of GSH, the transfection efficiency of SNC7 loaded with GFP mRNA in HEK293 and 3T3 cells were carried out under similar conditions as described above. However, in this case, GSH was intentionally added to the cell culture media with a GSH concentration ranging from 0.001 to 10 mM. Forty-eight hours after SNC treatment, the GFP expression level was measured using a flow cytometer and analyzed with FlowJo 7.6.

***In Vitro* CRISPR/Cas9 Genome Editing Efficiency Study**

For gene editing studies, GFP-HEK cells were used to test the CRISPR RNP and Cas9 mRNA/sgRNA delivery efficiency of SNCs *in vitro*. Cells were seeded at a density of 5,000 cells per well in a 96-well plate 24 h before treatment. RNP was complexed by mixing sNLS-*Sp*Cas9-sNLS and sgRNA at 1:1 in molar ratio for 5 min on ice. Cells were treated with RNP-encapsulated SNCs or RNP-complexed Lipo 2000 (0.5 μL/well). For each treatment, the RNP dosage was kept at 150 ng/well, with an equivalent Cas9 nuclease dosage at 125 ng/well. For mRNA:sgRNA weight ratio optimization, GFP-HEK cells were treated with SNCs encapsulating Cas9 mRNA (Trilink #L-7206) and sgRNA with different weight ratios ranging from 1:1 to 8:1 with the dose of total

RNA fixed at 200 ng/well. The gene editing efficiencies were quantified six days after treatment using flow cytometry by counting the percentage of green fluorescence negative cells. Data were analyzed with FlowJo 7.6.

The NIH 3T3 cells were used for *in vitro App* gene editing studies. Cells were seeded at a density of 5,000 cells per well in a 96-well plate 24 h before treatment. Cas9 mRNA and *App*$^{659}$ sgRNA were encapsulated into SNC with an mRNA : sgRNA weight ratio of 4:1 (i.e., mRNA/sgRNA-SNC). Other treatment groups include Lipo 2000 complexed with Cas9 mRNA and *App* sgRNA (i.e., mRNA/sgRNA-Lipo 2000) and RNP-encapsulated SNC (i.e., RNP-SNC). The dosage of the payload was 500 ng/well. The *in vitro* editing efficiency was analyzed 96 h post-treatment by next-generation sequencing (NGS).

**Subcellular Trafficking Study of the SNC**

Atto550-tagged RNP was used for the subcellular trafficking study of the SNC. The RNP was prepared as previously reported [3a, 4]. The guide RNA (i.e., gRNA) components (i.e., a negative control crRNA (Alt-R CRISPR-Cas9 Negative Control crRNA #1, Integrated DNA Technologies, Inc.) and an Atto550-tagged tracrRNA, 200 μM) were first mixed on ice with a 1:1 molar ratio, and then, sNLS–SpCas9–sNLS solution (62.5 μM) was added to the gRNA solution at a 1:1 molar ratio for 5 min on ice. The Atto550-tagged RNP was then encapsulated into SNC7.

Subcellular trafficking of the SNC loaded with Atto550-tagged RNP was studied by confocal laser scanning microscopy (CLSM, Nikon, Japan). Twenty-four hours before treatment, HEK 293 cells were seeded onto a Nunc Lab-Tek II CC2 Chamber Slide (Thermo Fisher, USA, 20,000 cells per well). At each time point (i.e., 0.5, 1, and 4 h) after SNC treatment, the live cells were stained with endosome/lysosome marker LysoTracker Green DND-26 (100 nM) for 30 min at 37 °C, and then

Hoechst33342 (5 μg/ml) for 5 min at room temperature. The subcellular localization of Atto550-tagged RNP was analyzed by CLSM.

**Cell Viability Assay**

The cytotoxicity of SNCs was studied by an MTT assay. Cells were treated with complete medium, DNA-complexed Lipo 2000 (0.5 μL/well), and DNA-loaded SNCs. Cell viability was measured using a standard MTT assay 48 h after treatment (Thermo Fisher, USA). Briefly, cells were treated with media containing 500 μg/mL MTT and incubated for 4 h. Then, the MTT-containing media was aspirated, and the purple precipitate was dissolved in 150 μL of DMSO. The absorbance at 560 nm was obtained with a microplate reader (GloMax® Multi Detection System, Promega, USA).

**Animal**

All animal experiments were conducted in strict adherence to the Guide for the Care and Use of Laboratory Animals (National Institutes of Health). Animal protocols were approved by the Institutional Animal Care and Use Committee (IACUC) at the University of Wisconsin-Madison (Protocol ID: M006286). B6.Cg-Gt(ROSA)26Sor$^{tm14(CAG-tdTomato)Hze}$/J mice (Ai14 mice, 6 to 8 weeks old, male and female, JAX #007914) were used for *in vivo* Cre mRNA and RNP delivery experiments. C57BL/6J (wild-type mice, 6 to 9 weeks old, JAX #000664) were used for biodistribution studies, as well as *App*- and *Th*- gene editing experiments. All mice were maintained under a tightly controlled temperature (22 °C), humidity (40–50%), and light/dark (12/12 h) cycle conditions, with *ad libitum* access to water and food. All mice were randomly divided into groups, prior to injection/processing. Mice were anesthetized with 5% isoflurane for 5-10 min, then maintained in 1% isoflurane during injection/processing.

***In Vivo* Cre mRNA and RNP Delivery in Ai14 Mice**

Ai14 mice were used to assess the Cre mRNA transfection/editing efficiency and RNP editing efficiency induced by Cre mRNA- or Cas9/sgRNA RNP-encapsulated SNCs, respectively. Cre-mRNA was purchased from Trilink Biotechnologies, USA (#L-7211). RNPs were prepared using a sgRNA targeting the stop cassette composed of 3× SV40 polyA blocks. Ai14 mice were subjected to fasting for 24 h and injected with SNCs through retro-orbital injections (injection volume: 100 μL, dosages of the payload were specified). PBS-injected Ai14 mice were used as controls. Thirty minutes post-injection, blood glucose was restored by intraperitoneal injection of 200 μl of 20 wt% D-(+)-glucose solution in PBS. For single injection, the SNC-injected and control mice were perfused with ice-cold PBS 14 days post-injection. For triple injection (Fig. 4D), mice were injected 3 times at 5-day intervals and sacrificed 15 days after the first injection. Organs and tissues (brain, liver, heart, lungs, spleen, kidneys, and muscle) were then collected and analyzed. Fresh organs/tissues were imaged using the *in vivo* imaging system (IVIS Lumina system, Perkin Elmer) for tdTomato expression, and then processed for immunofluorescence flow cytometry (IFCM) and CLSM analyses.

**Biodistribution Study**

Wild-type C57BL/6J mice (n = 3) were subjected to fasting for 24 h. These mice were retro-orbitally injected with 100 μl of SNC7-10%RVG+10%Glu encapsulating Atto550-RNP. After 30 min, these mice were intraperitoneally injected with 200 μl of 20 wt% D-(+)-glucose solution in PBS. In another group, free-feeding C57BL/6J mice (n=3) were injected with the same SNC formulation without glycemic control. Twenty-four hours post-SNC injection, mice were perfused with ice-cold PBS, and major organs including the brain, liver, lungs, kidneys, spleen, heart, and muscle were collected and washed with PBS. Thereafter, a piece of the tissue sample was accurately weighed after removing excess fluid and homogenized with cell lysis buffer (i.e., RIPA buffer supplemented with cOmplete Protease Inhibitor Tablets (Thermo Fisher)). After the homogenized tissue solution was centrifuged at 12,000 × g for 30 min at 4 °C, the supernatant was collected and the Atto550 fluorescence intensity was quantified using a microplate reader (GloMax Multi Detection System, Promega, USA).

***In Vivo App* and *Th* Gene Editing**

*App*-SNC was prepared by co-encapsulating Cas9 mRNA and $App^{659}$ sgRNA into SNC7-10%RVG+10%Glu with a 4:1 weight ratio. *Th*-SNC was prepared similarly using a *Th*-targeting sgRNA. The protospacer sequences of $App^{659}$ sgRNA and *Th*-targeting sgRNA can be found in Table S1.

Wild-type C57BL/6J mice (n=7) were retro-orbitally injected with *App*-SNC under glycemic control following the triple injection timeline (Fig. 4D). For each injection with glycemic control, mice were subjected to fasting for 24 h. And then, these mice were retro-orbitally injected with 100 µl of SNC7-10%RVG+10%Glu with the corresponding payloads. After 30 min, these mice were intraperitoneally injected with 200 µl of 20 wt% D-(+)-glucose solution in PBS. PBS-injected mice were used as controls. Fifteen days after the first injection, mice were separated into two subgroups randomly, one subgroup (n=4) was anesthetized, and the whole blood was collected from the sublingual vessel into heparin-coated collection tubes and the serum was separated by centrifugation at 1500 × g for 10 min for hematological analysis. Thereafter, mice were euthanized, and their major organs and tissues were collected freshly for NGS, western blot, and RT-qPCR analyses. The other subgroup (n=3) was perfused with ice-cold PBS and organs were fixed for CLSM imaging.

**Brain Cell Collection for IFCM**

The brain cells were dissociated and collected following previously established protocols with minor changes [5]. Briefly, after PBS perfusion and tissue collection, brains were cut into 1 mm pieces coronally using a coronal mouse brain matrix (CellPoint Scientific, USA) on ice. The brain pieces containing the cortex, hippocampus and thalamus/hypothalamus were mixed with 50 μL Hibernate A low fluorescence buffer (Brain Bits, USA) and thoroughly minced on a glass slide on the ice. The minced tissues were transferred into 1.5 mL Hibernate A low fluorescence buffer and precipitated by centrifugation at 200 × g for 4 min at 4 °C. The pellet was then mixed with 1.5 mL cold Accutase cell detachment solution (ThermoFisher, USA) for 30 min at 4 °C before centrifugation and resuspension in Hibernate A low fluorescence buffer. The digested tissue was mechanically triturated and fixed with 50% ethanol at 4 °C. The cell suspension was re-dispersed in Hibernate A low fluorescence buffer and filtered through cell strainers (100 μm and 40 μm) before immunofluorescence staining.

**Immunofluorescence Staining for IFCM and CLSM Analyses**

For IFCM analysis, brain cells were incubated in 10% goat serum and 0.3% Triton X-100 in PBS at RT for 1h. The cell samples were then separated into 3 equal aliquots used for cell marker and RFP primary antibodies (1 h) and their corresponding secondary antibodies (1 h) (Table S2 and S3). After antibody staining, the cell samples were then stained with DAPI and resuspended in cold PBS for storage and IFCM analysis.

For CLSM imaging, tissues were fixed in 4% paraformaldehyde (PFA) at 4 °C for 48 h followed by incubation in a PBS solution containing 30% sucrose and stored at 4 °C for 72 h. Thereafter, the fixed and dehydrated tissues were embedded in Tissue-Tek Optimal Cutting Temperature (OCT) Compound (Sakura Finetek, USA) and then frozen in dry ice. The blocks were sectioned using a cryostat machine (CM1900, Leica Biosystems, USA) at 50 μm thickness and mounted on microscope slides. The sections were incubated in 10% goat serum and 0.3% Triton X-100 in PBS for 1h. For immunofluorescence staining, the sections were first incubated with corresponding primary antibodies for 1 h at room temperature (Table S2). The primary antibody was then recognized by a fluorescence-conjugated secondary antibody (Table S3). Finally, the slides were stained with DAPI, mounted with ProLong Gold Antifade Reagent, and covered with microscope cover glasses. All the images were acquired using a Nikon AR1 confocal laser scanning microscope (CLSM).

**Next-Generation Sequencing**

The genomic DNA from 3T3 cells, brain regions, and other organs was extracted using Monarch Genomic DNA Purification Kit (New England Biolabs, #T3010) following the manufacturer's protocol. Genomic PCR was performed using Q5 High-Fidelity 2X Master Mix (New England Biolabs) with genomic DNA templates (50 ng) and customized primers ($0.5 \times 10^{-6}$ M, Integrated DNA Technologies, Inc., as listed in Table S4), following the manufacturer's protocol. PCR products were purified using AMPure XP (Beckman Coulter) and quantified by Nanodrop One Microvolume UV–vis Spectrophotometer (Thermo Fisher Scientific). Individual samples were pooled and run on an Illumina MiniSeq Sequencing System at a run length of 2 × 150 bp. Editing efficiency was analyzed by CRISPResso2 [6].

**Western Blot**

Western blot (WB) was used to analyze the level of intact APP and TH in the brain tissue. Cell lysates were prepared in cell lysis buffer. After the cell lysates were centrifuged at 12,000 × g for 15 min at 4 °C, supernatants were quantified and resolved by SDS-PAGE for western blot analysis. The APP Y188 antibody was used to detect the intact APP in the cell lysate, while a tyrosine hydroxylase antibody (EP1532Y) was used to detect TH in the cell lysate (Table S2 and S3).

Glyceraldehyde 3-phosphate dehydrogenase (GAPDH) was used as a loading control (antibodies and dilution factors are listed in Table S2 and S3). Band intensities were analyzed by Empiria Studio Software (LI-COR).

***In Vivo* Biocompatibility Assay**

Hematological analysis was performed using the collected mouse serum to evaluate the key elements of the blood biochemical profile using VetScan Preventive Care Profile Plus rotors (Abaxis) in a VetScan VS2 blood chemistry analyzer (Abaxis), following the manufacturer's protocol. To further evaluate the systemic or local toxicity, tissue sections were stained with Hematoxylin and eosin (H&E) and observed under an optical microscope.

***In Vivo* Immunogenicity Assay**

The immunogenicity of SNC was analyzed by RT-qPCR. In brief, a portion of the brain tissue was collected and stored in RNAlater solution (Thermo Fisher Scientific) at -20 °C until RNA extraction. RNA in these samples was extracted using TRIzol reagent (Thermo Fisher Scientific) following the manufacturer's protocol and quantified by Nanodrop One Microvolume UV-vis Spectrophotometer. The cDNA was then synthesized using iScript Reverse Transcription Supermix (Bio-Rad Laboratories, Inc.) following the manufacturer's protocol. RT-qPCR was finally performed using iTaq Universal SYBR Green Supermix (Bio-Rad Laboratories, Inc.) with the cDNA templates (10 ng) and customized primers ($0.5 \times 10^{-6}$ M, Integrated DNA Technologies, Inc.) for genes of interest (Table S4) on a CFX96 Touch Real-Time PCR Detection System (Bio-Rad Laboratories, Inc.). GAPDH was selected as the reference gene for data analysis. The thermocycling for qPCR was: 40 cycles of 95 °C for 5 s and 60 °C for 30 s. Melt curve analysis was performed at the end of qPCR experiments. Data were analyzed using CFX Maestro 2.0 (Bio-Rad).

**Statistical Analysis**

Results are presented as mean ± standard deviation (SD). One-way analysis of variance (ANOVA) with Tukey's multiple comparisons was used to determine the difference between independent groups. Statistical analyses were conducted using GraphPad Prism 8 software.

### Supplementary Figures

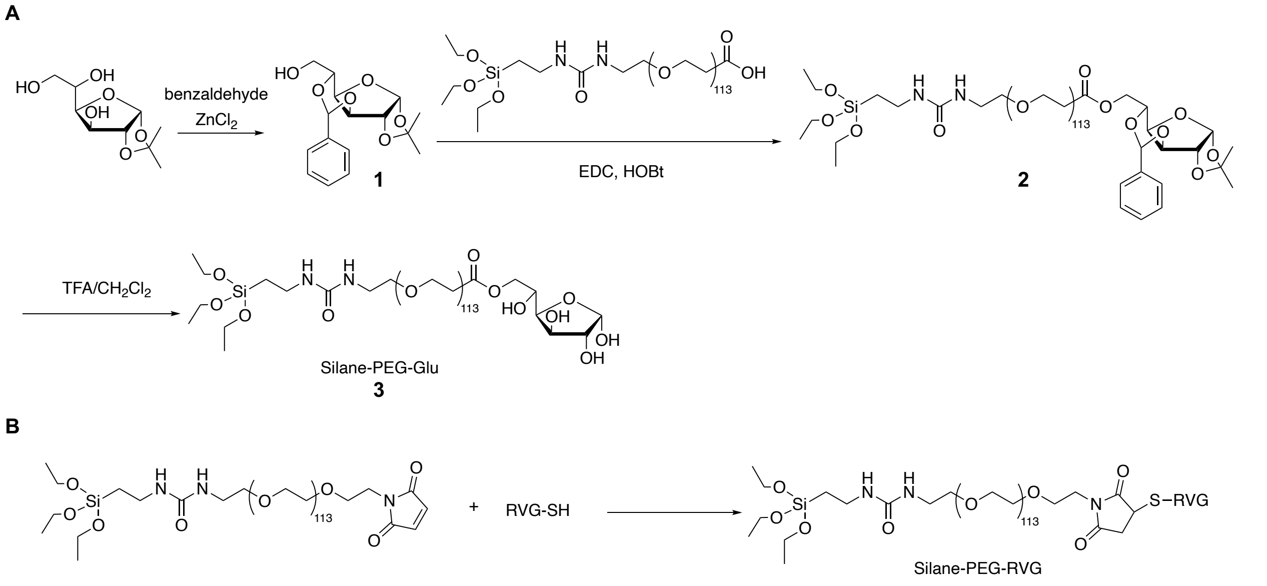


**Figure S1**. The synthesis scheme of PEG-silane conjugated with different targeting ligands. **(A)** silane-PEG-Glu, and **(B)** silane-PEG-RVG. Glucose (Glu) was conjugated onto silane-PEG-COOH to obtain silane-PEG-Glu via an esterification reaction between the carboxyl group on PEG and the hydroxyl group at the C6 position of glucose. Silane-PEG-RVG was synthesized via a thiol-maleimide Michael addition reaction.

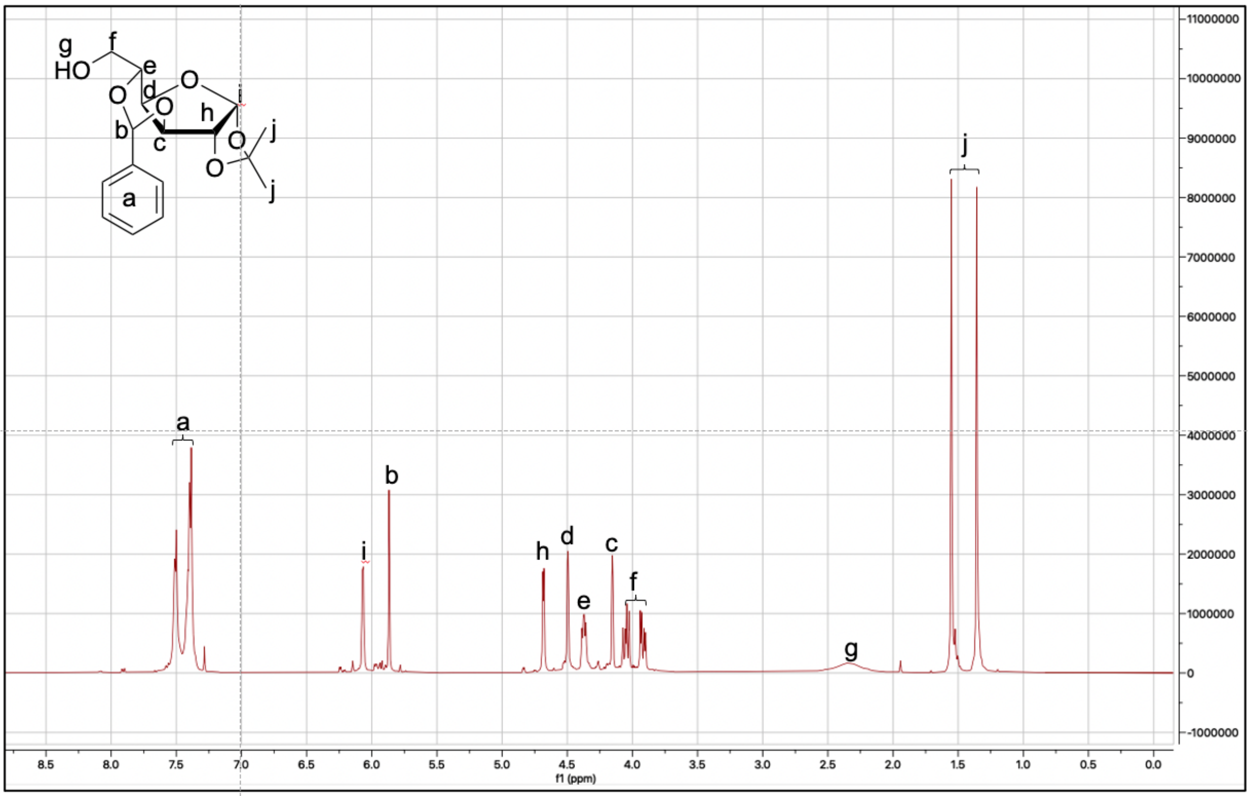


**Figure S2**. $^{1}$H NMR spectrum of 3,5-*O*-benzylidene-1,2-*O*-isopropylidene-*α*-*D*-glucofuranoside (BIG).

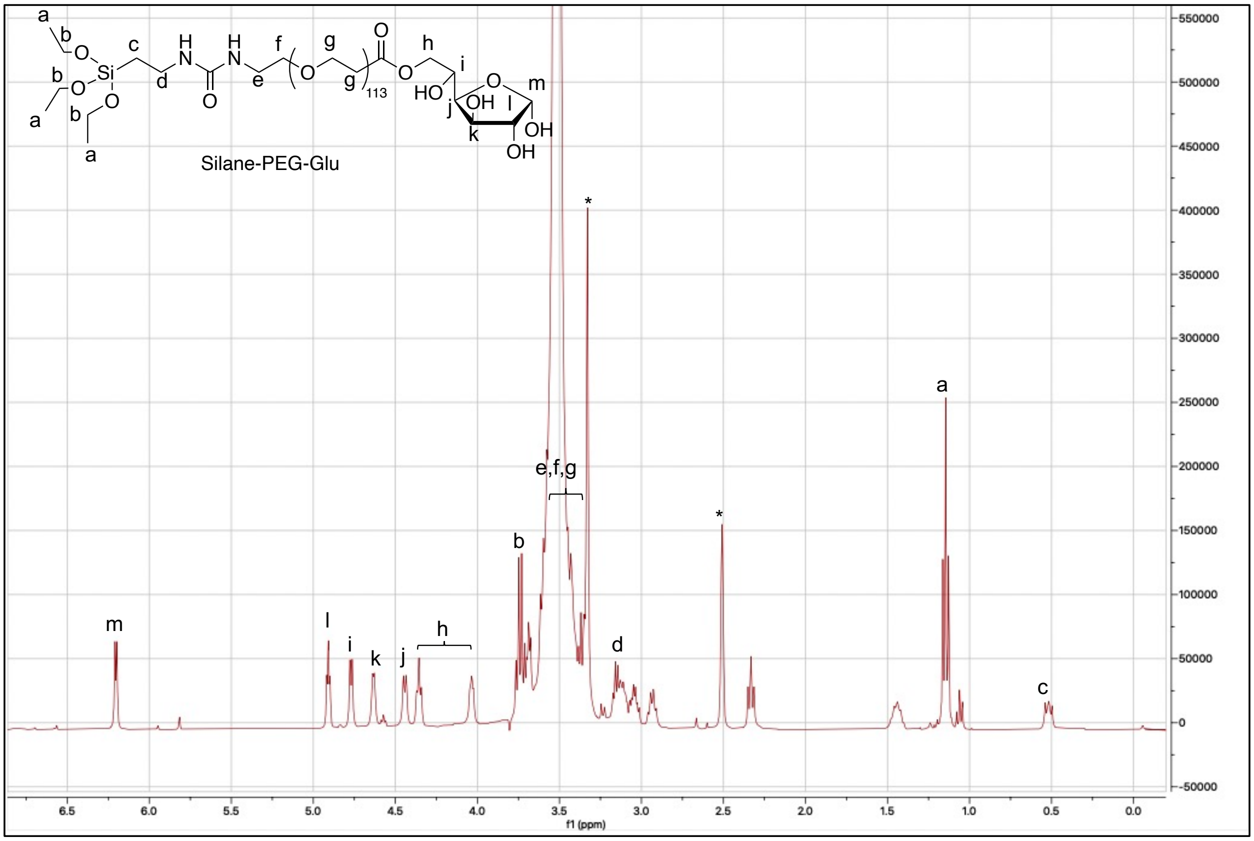


**Figure S3**. $^{1}H$ NMR spectrum of silane-PEG-Glu.

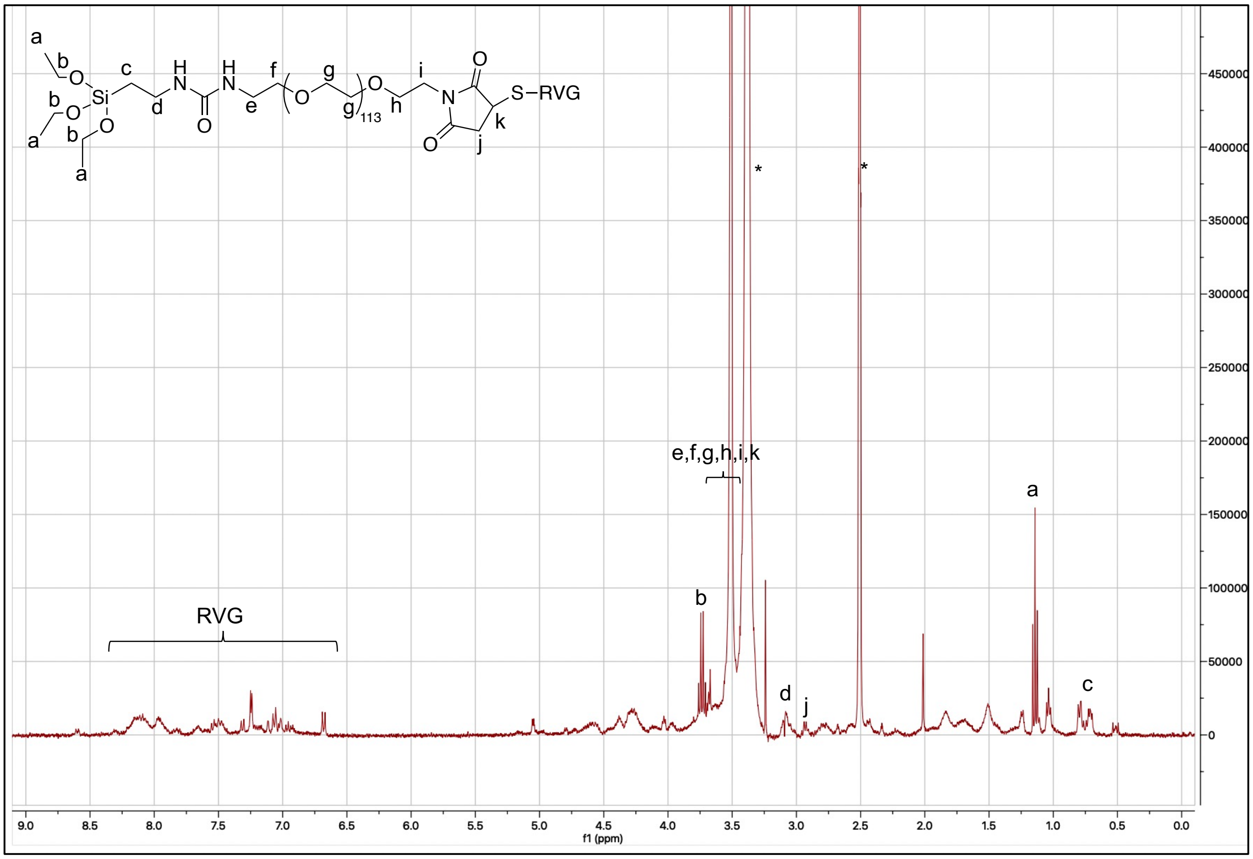


**Figure S4**. $^{1}$H NMR spectrum of silane-PEG-RVG.

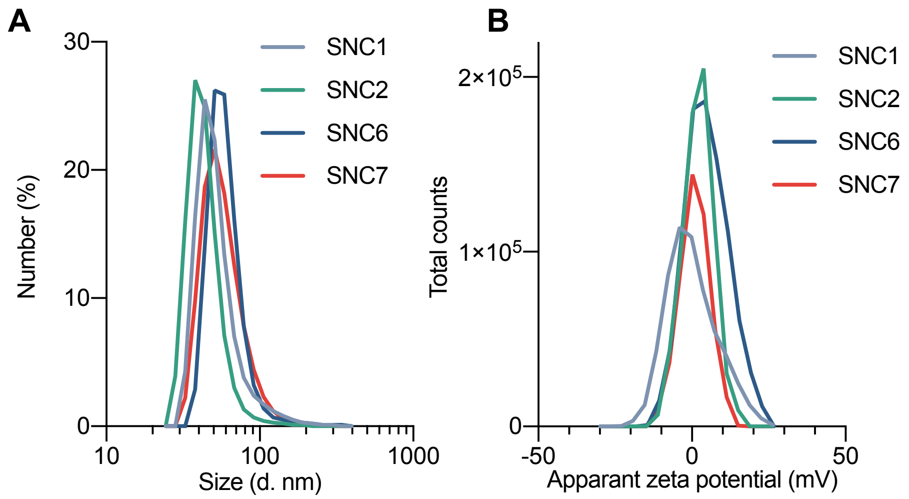


**Figure S5. (A)** Size distribution of DNA-encapsulated SNC1, SNC2, SNC6 and SNC7 measured by DLS. **(B)** Zeta-potentials of DNA-encapsulated SNC1, SNC2, SNC6 and SNC7.

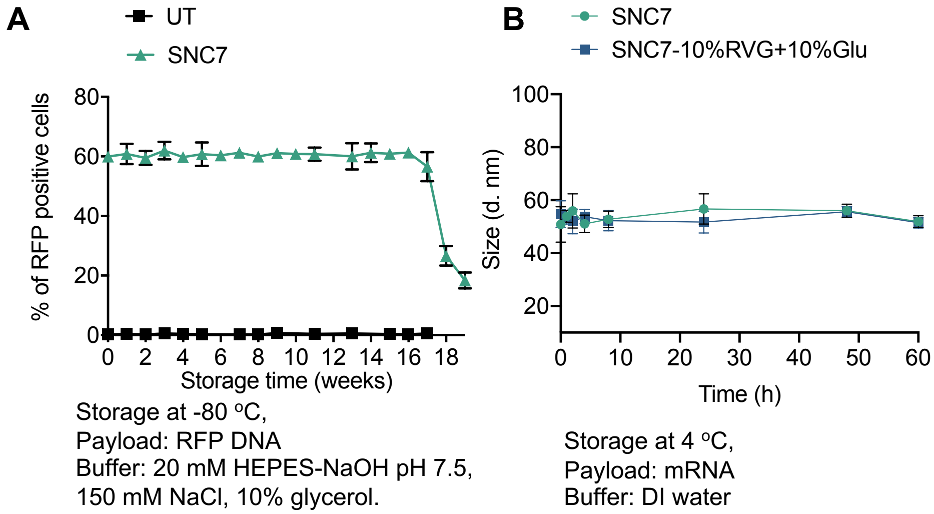


**Figure S6**. Stability studies of SNC7. **(A)** DNA transfection efficiency of SNC7 after storage at -80 °C in a storage buffer (i.e., 20 mM HEPES-NaOH pH 7.5, 150 mM NaCl, 10% glycerol). UT: untreated. **(B)** Size of SNC7 with or without targeting ligands in aqueous solution after storage at 4 °C.

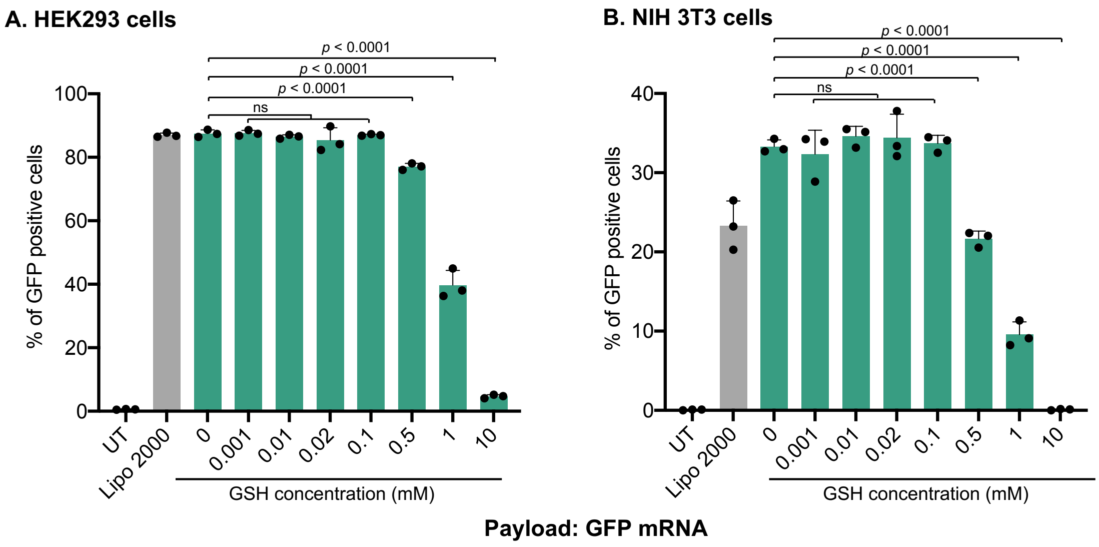


**Figure S7**. The effect of GSH concentration in cell culture medium on the mRNA transfection efficiency of SNC7 in HEK293 **(A)** and NIH 3T3 **(B)** cells. Data are presented as mean ± SD. Statistical significance was calculated with PBS-injected as the control via one-way ANOVA with Tukey's post hoc test. ns, not significant.

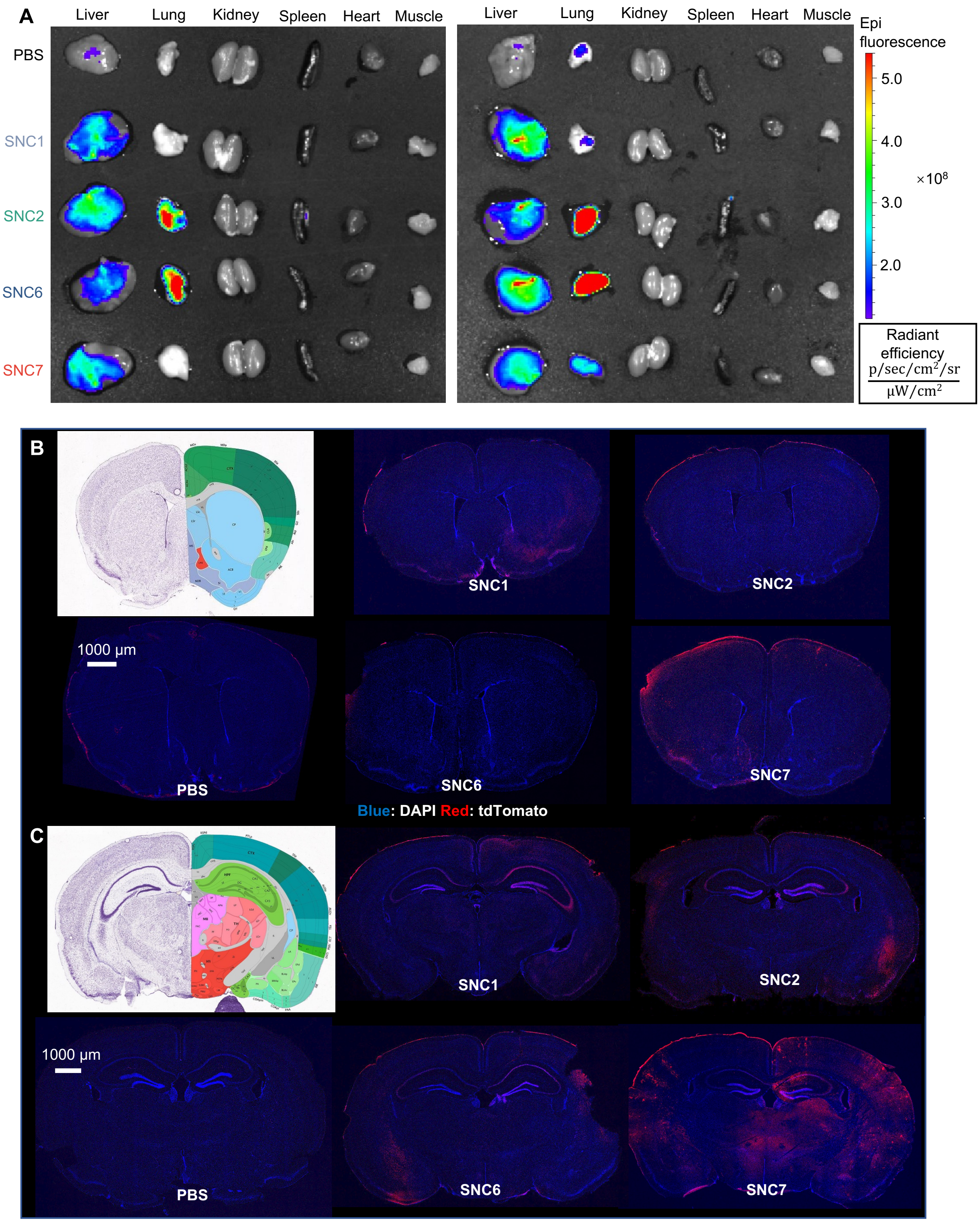
A
Liver
Lung
Kidney
Spleen
Heart
Muscle
PBS
SNC1
SNC2
SNC6
SNC7
Epi fluorescence
5.0
4.0
3.0
2.0
×10^8
Radiant efficiency
p/sec/cm²/sr
μW/cm²
B
SNC1
SNC2
1000 μm
PBS
SNC6
SNC7
Blue: DAPI Red: tdTomato
C
SNC1
SNC2
1000 μm
PBS
SNC6
SNC7

**Figure S8. (A)** TdTomato signal in major organs of Ai14 mice intravenously injected with Cre mRNA-encapsulated SNCs of different formulations (SNC1, SNC2, SNC6 and SNC7) with the same targeting ligand combination (10%RVG+10%Glu). **(B-C)** Representative coronal section mosaic tile CLSM images of the brains of Ai14 mice intravenously injected with Cre mRNA-encapsulated SNCs of different formulations at +1.18 mm bregma **(B)** and -1.70 mm bregma (Cre mRNA dosage: 2 mg/kg)**(C)**.

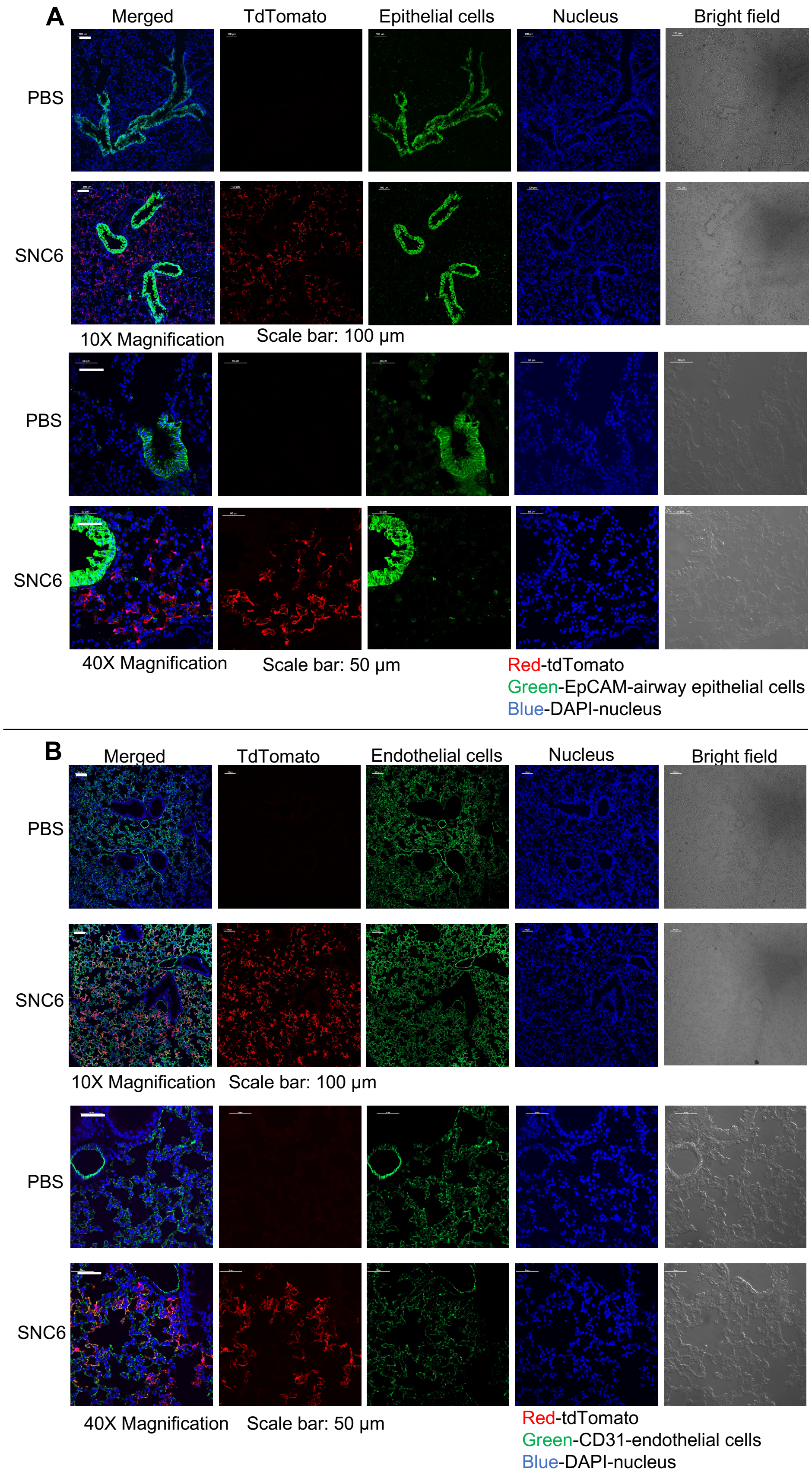

A
Merged
TdTomato
Epithelial cells
Nucleus
Bright field
PBS
SNC6
10X Magnification
Scale bar: 100 μm
PBS
SNC6
40X Magnification
Scale bar: 50 μm
Red-tdTomato
Green-EpCAM-airway epithelial cells
Blue-DAPI-nucleus
B
Merged
TdTomato
Endothelial cells
Nucleus
Bright field
PBS
SNC6
10X Magnification
Scale bar: 100 μm
PBS
SNC6
40X Magnification
Scale bar: 50 μm
Red-tdTomato
Green-CD31-endothelial cells
Blue-DAPI-nucleus

**Figure S9.** Intravenous injection of Cre mRNA-encapsulated SNC6-10%Glu+10%RVG induced tdTomato expression in the lungs of Ai14 mice. Major cell types with tdTomato expression are pulmonary vascular endothelial cells and/or alveolar epithelial cells, but not airway epithelial cells. **(A)** Representatives CLSM image of lung slices to identify tdTomato positive cells and EpCAM (an airway epithelial cell marker) positive cells by immunofluorescence staining. **(B)** Representatives CLSM image of lung slices immunostained to identify tdTomato positive cells co-localizing with CD31 (a capillary endothelial cell marker) positive cells.

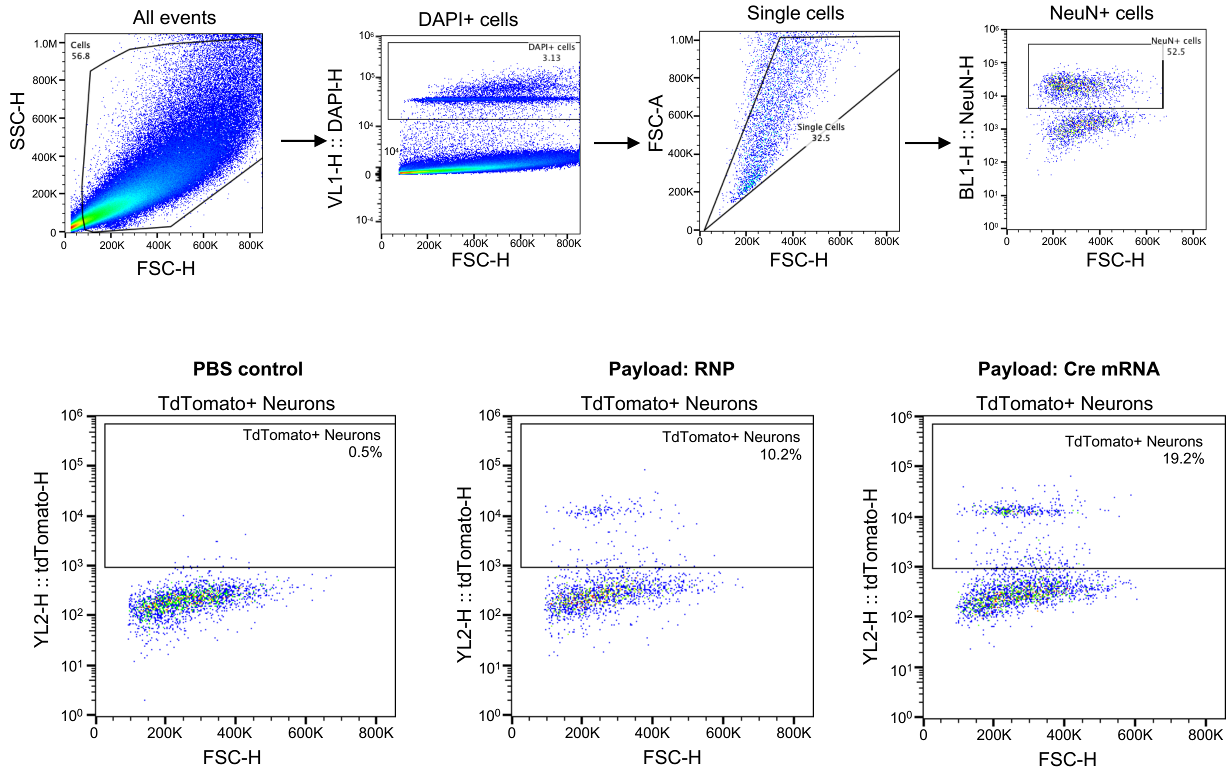


**Figure S10.** Gating strategy of tdTomato positive neurons. The 'Cells' gate was determined by forward and side scatter properties and confirmed by positive labeling with nuclear DAPI staining. The 'NeuN+ cells' gate in the cell population was determined by the fluorescence of NeuN, a neuron marker. TdTomato+ cells were identified in the 'NeuN+ cells' gate by tdTomato fluorescence.

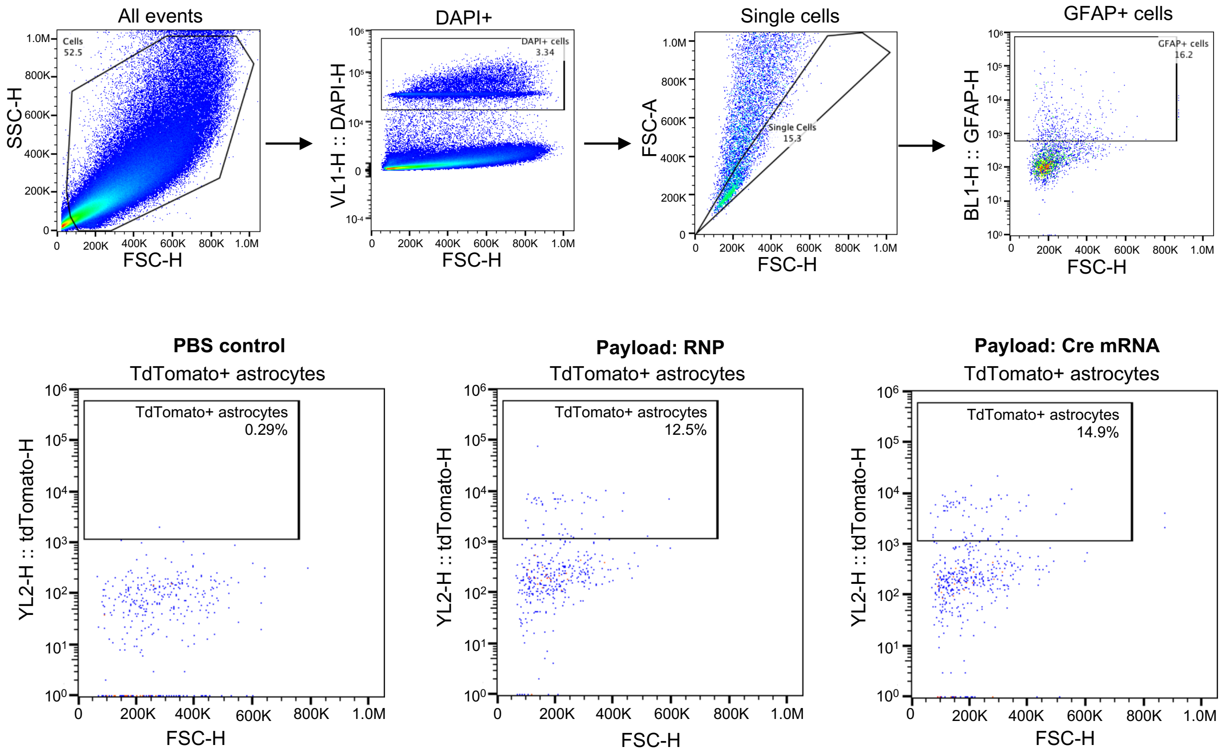


**Figure S11.** Gating strategy of tdTomato positive astrocytes. The 'Cells' gate was determined by forward and side scatter properties and confirmed by positive labeling with nuclear DAPI staining. The 'GFAP+ cells' gate in the cell population was determined by the fluorescence of GFAP, an astrocyte marker. TdTomato+ cells were identified in the 'GFAP+ cells' gate by tdTomato fluorescence.

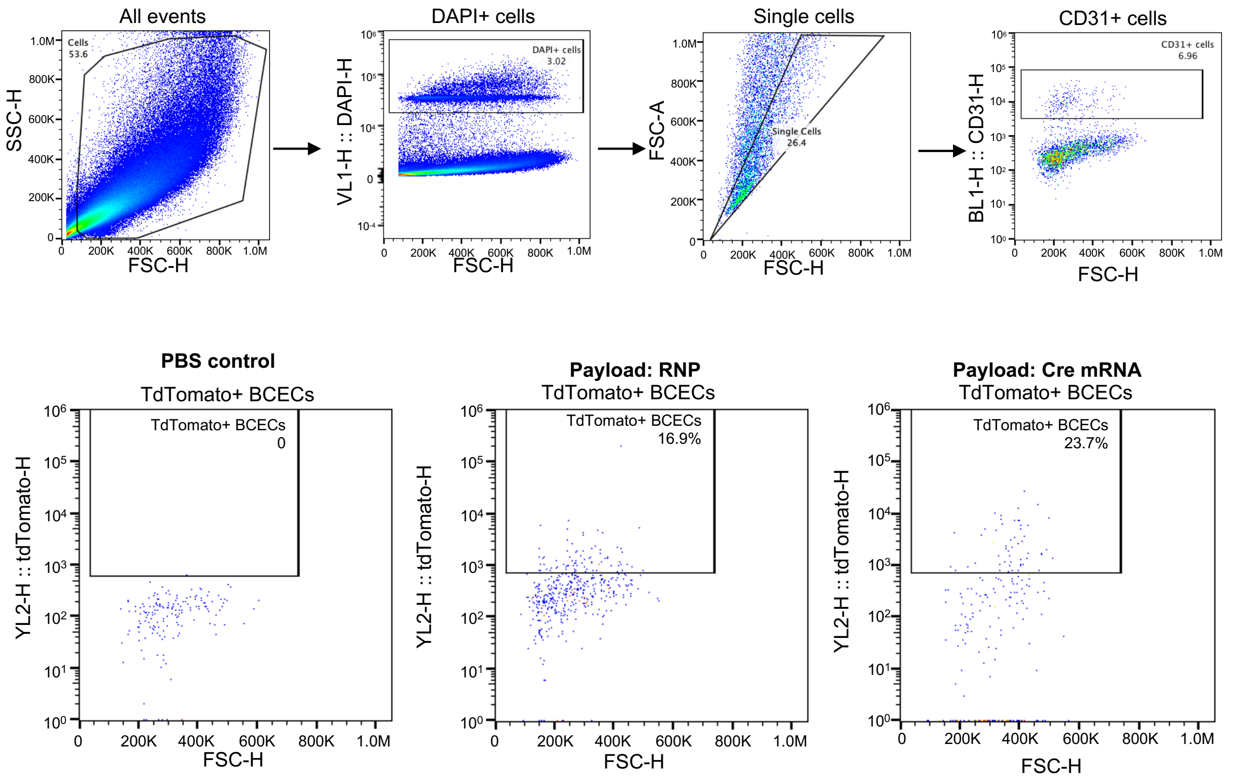


**Figure S12.** Gating strategy of tdTomato positive BCECs. The 'Cells' gate was determined by forward and side scatter properties and confirmed by positive labeling with nuclear DAPI staining. The 'CD31+ cells' gate in the cell population was determined by the fluorescence of CD31, a BCEC marker. TdTomato+ cells were identified in the 'CD31+ cells' gate by tdTomato fluorescence.

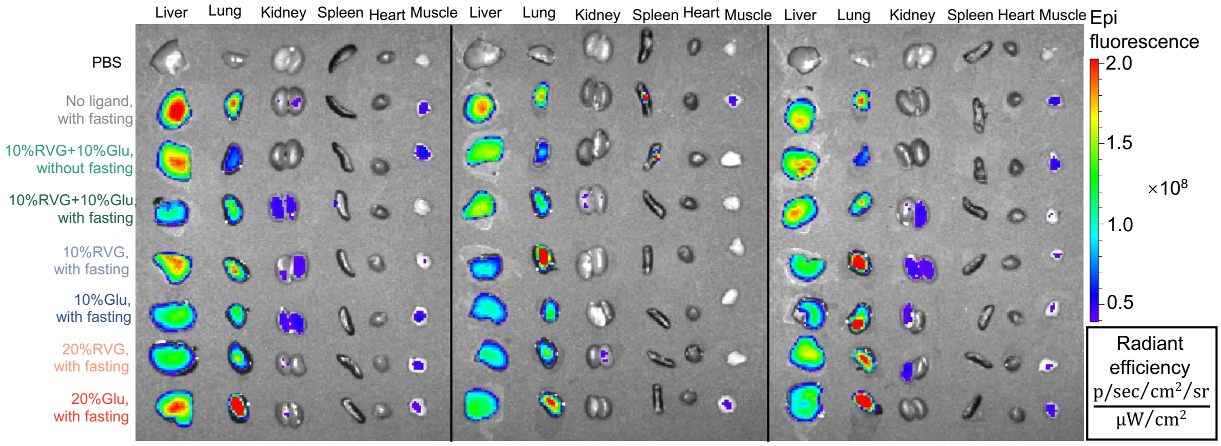


**Figure S13.** TdTomato signal in the major organs of Ai14 mice intravenously injected with RNP-encapsulated SNC7 conjugated with different amounts of glucose and RVG peptide with or without glycemic control (n=3).

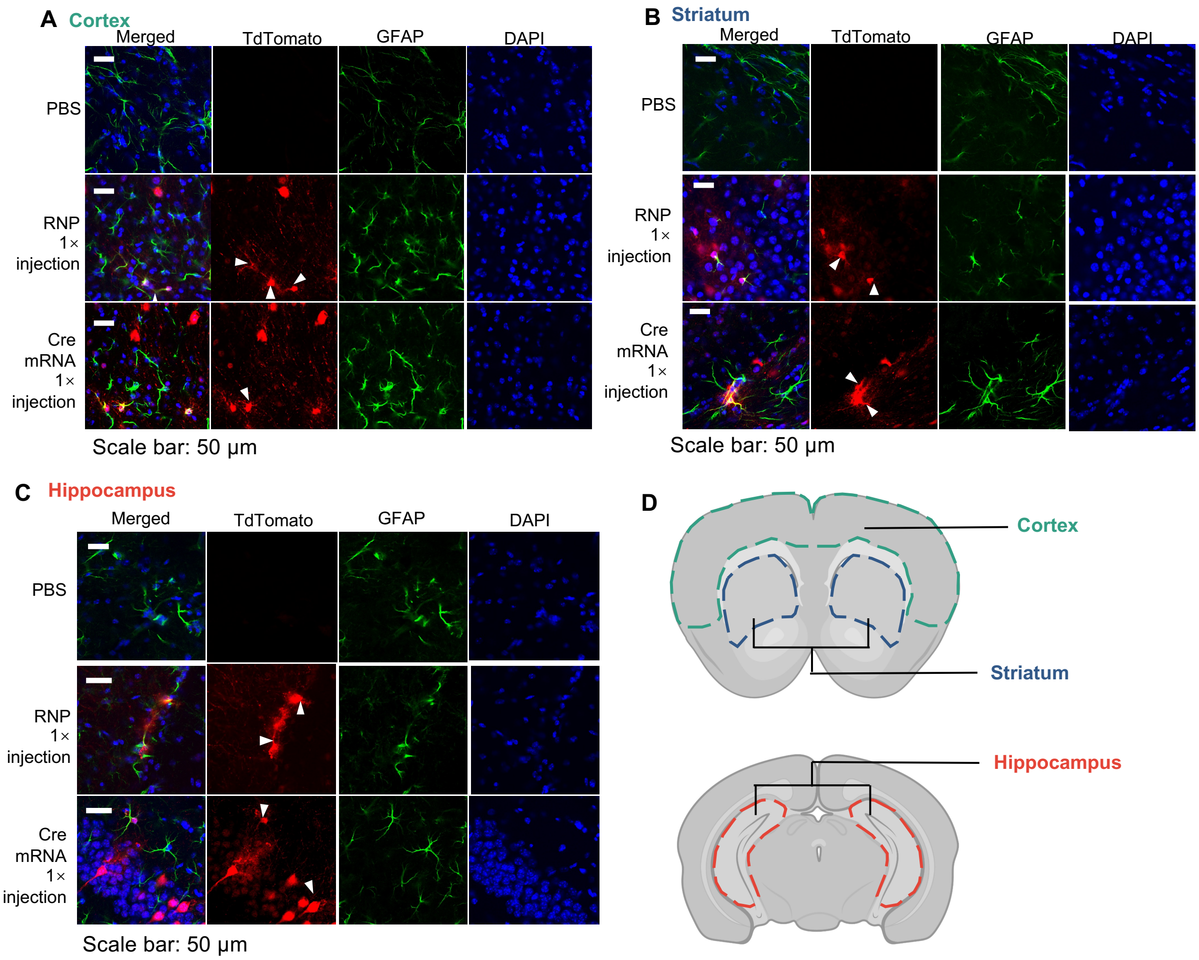


**Figure S14.** Intravenous injection of RNP- or Cre mRNA-encapsulated SNCs induced tdTomato expression in astrocytes of Ai14 mice in **(A)** cortex, **(B)** striatum, and **(C)** hippocampus. TdTomato (red), nuclei (blue) and Astrocytes (GFAP+, green) were stained by corresponding antibodies. White arrowheads are pointing to tdTomato-positive astrocytes. The formulation used in this study was SNC7-10%RVG+10%Glu. **(D)** Regions of interest for CLSM imaging. Colored dash lines enclose the cortex, striatum and hippocampus.

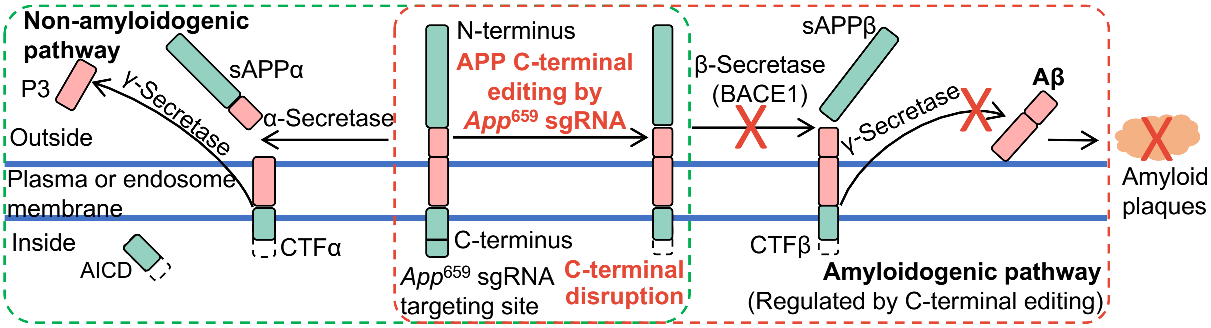


**Figure S15.** Schematic illustration of APP processing pathways (i.e., non-amyloidogenic and amyloidogenic). Editing the C-terminus of APP using *App*$^{659}$ sgRNA down-regulates APP-β-cleavage and Aβ formation. CTF: C-terminal fragment; AICD: APP intracellular domain.

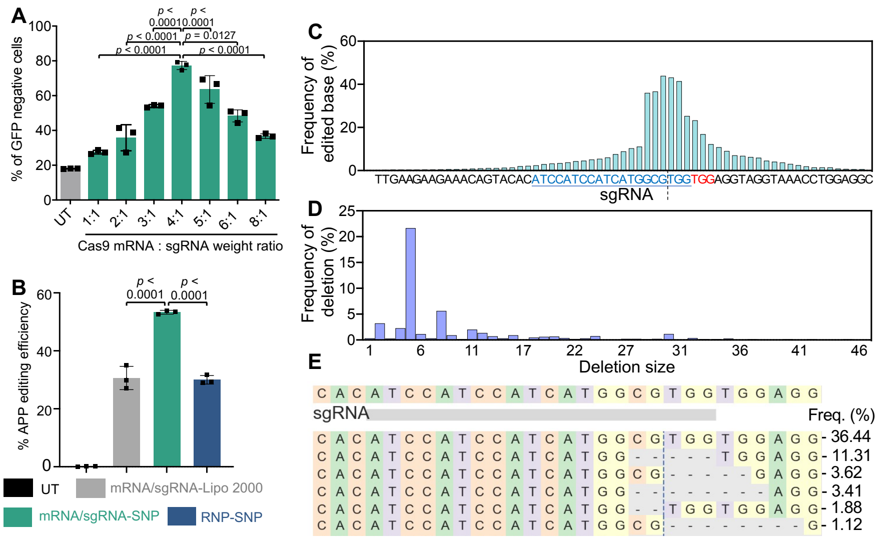


**Figure S16. (A)** Optimization of Cas9 mRNA:sgRNA weight ratio in GFP-HEK cells. Cas9 mRNA and sgRNA targeting GFP were encapsulated into SNC with different weight ratios ranging from 1:1 to 8:1, and treated with GFP-HEK cells (in a 96-well plate) with a total RNA dose of 200 ng/well (n=3). The Cas9 mRNA: sgRNA weight ratio of 4:1 induced the highest gene editing efficiency. **(B)** The *App* gene editing efficiency of SNC without any ligands was studied in NIH 3T3 fibroblasts. Cas9 mRNA and *App*$^{659}$ sgRNA were encapsulated into SNC with an mRNA: sgRNA weight ratio of 4:1 (i.e., mRNA/sgRNA-SNC). Other treatment groups include Lipo 2000 complexed with Cas9 mRNA and *App*$^{659}$ sgRNA (i.e., mRNA/sgRNA-Lipo 2000) and RNP-encapsulated SNC (i.e., RNP-SNC). The *in vitro* editing efficiency (n=3) was analyzed 96 h post-treatment by next-generation sequencing (NGS). Among the three treatment groups, mRNA/sgRNA-SNC exhibited significantly higher *App* gene editing efficiency (54%) than mRNA/sgRNA-Lipo2000 (31%) and RNP-SNC (30%). Thus, mRNA/sgRNA-SNC was used for *in vivo* studies. **(C)** Representative spectrum showing the frequency of editing for every base proximal to the sgRNA targeting sequence. **(D)** Representative spectrum showing the size of deletions in the edited sequencing reads. **(E)** Frequencies of major reads with editing. Data are presented as mean ± SD. Statistical significance was calculated with PBS-injected as the control via one-way ANOVA with Tukey's post hoc test. ns, not significant.

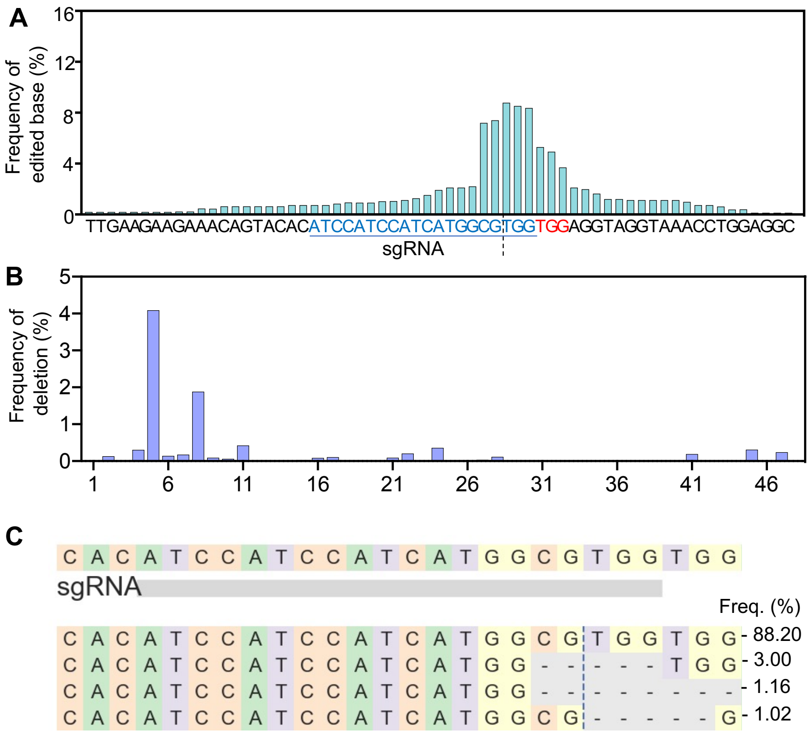


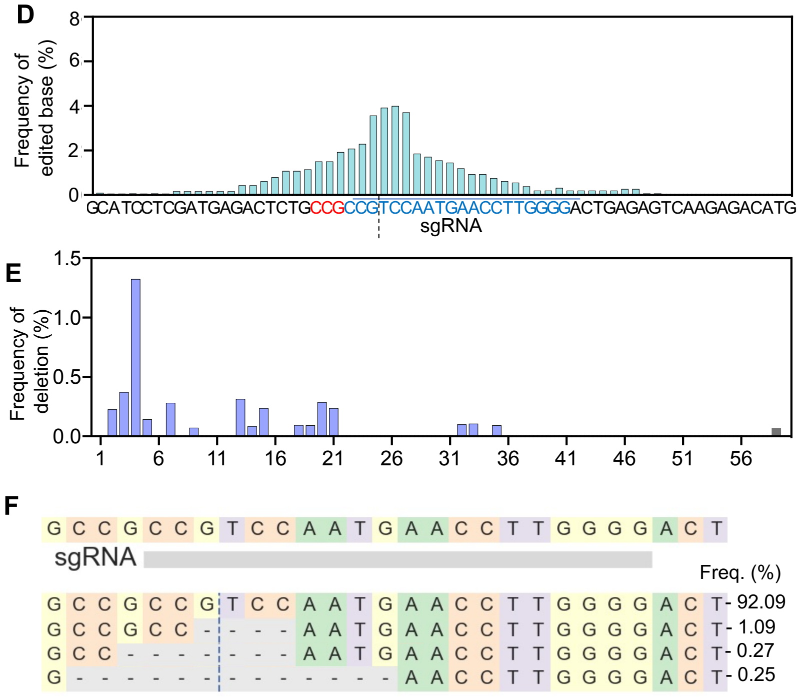

**Figure S17.** **(A-C)** Supplementary NGS results for *App* gene editing by *App*-SNC in wild-type mice. **(A)** Representative spectrum showing the frequency of editing for every base proximal to the sgRNA targeting sequence. **(B)** Representative editing spectrum showing the size of deletions in the edited sequencing reads. **(C)** Frequencies of major reads after *App* gene editing. **(D-F)** Supplementary NGS result for *Th* gene editing by *Th*-SNC in wild-type mice. **(D)** Representative spectrum showing the frequency of editing for every base proximal to the sgRNA targeting sequence. **(E)** Representative editing spectrum showing the size of deletions in the edited sequencing reads. **(F)** Frequencies of major reads after *Th* gene editing.

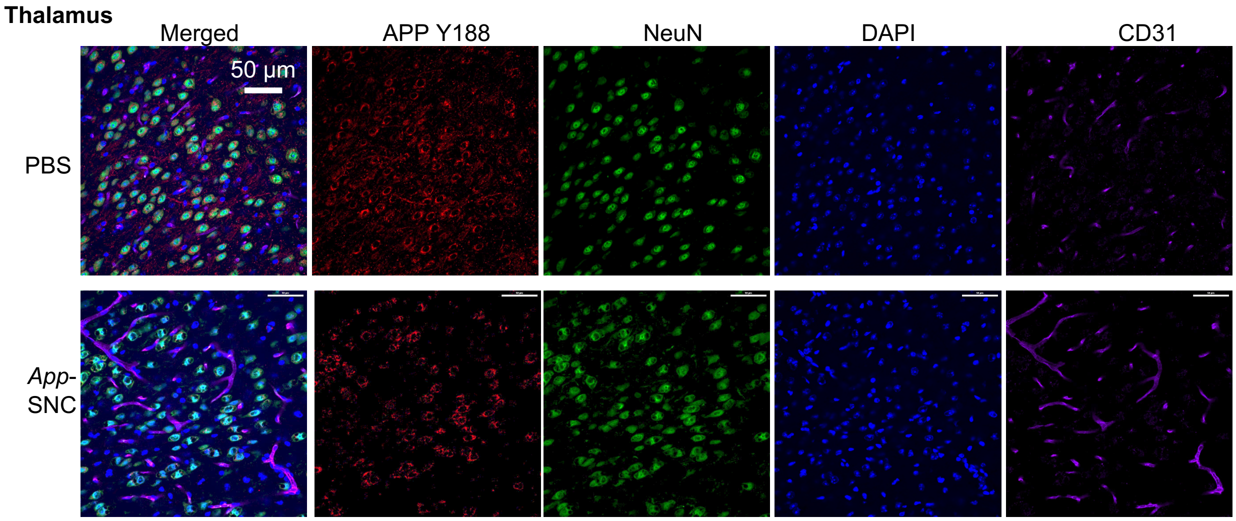


**Figure S18.** CLSM images of mouse thalamus with intravenous injection of *App*-SNC or PBS (control). Brain slices were stained with antibodies against APP C-terminus (APP-Y188) (red), NeuN (green), CD31 (magenta), and DAPI (blue). Attenuation of Y-188 staining indicates successful APP C-terminal editing.

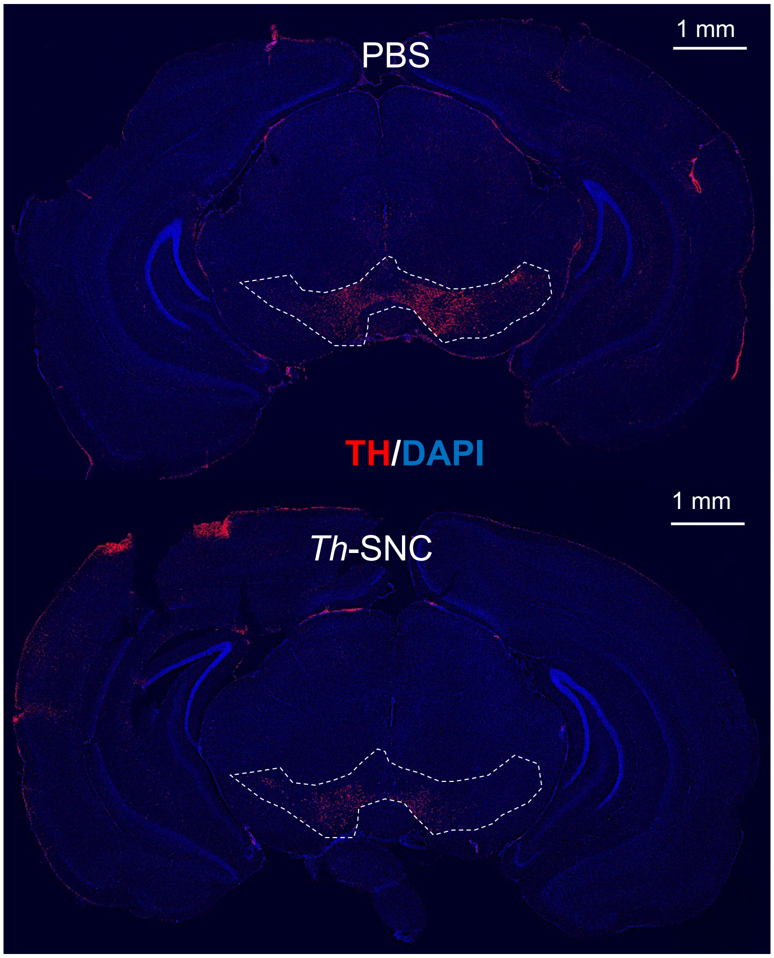


**Figure S19.** Representative coronal section mosaic tile CLSM images of the brains (at -3.50 mm bregma) of C57BL/6 mice intravenously injected with *Th*-SNC. Brain slices were stained with anti-TH antibody and DAPI. Major TH-expressing regions of the brain (i.e., ventral tegmental area and substantia nigra) are enclosed by white dash lines. Red: TH, blue: DAPI staining nuclei.

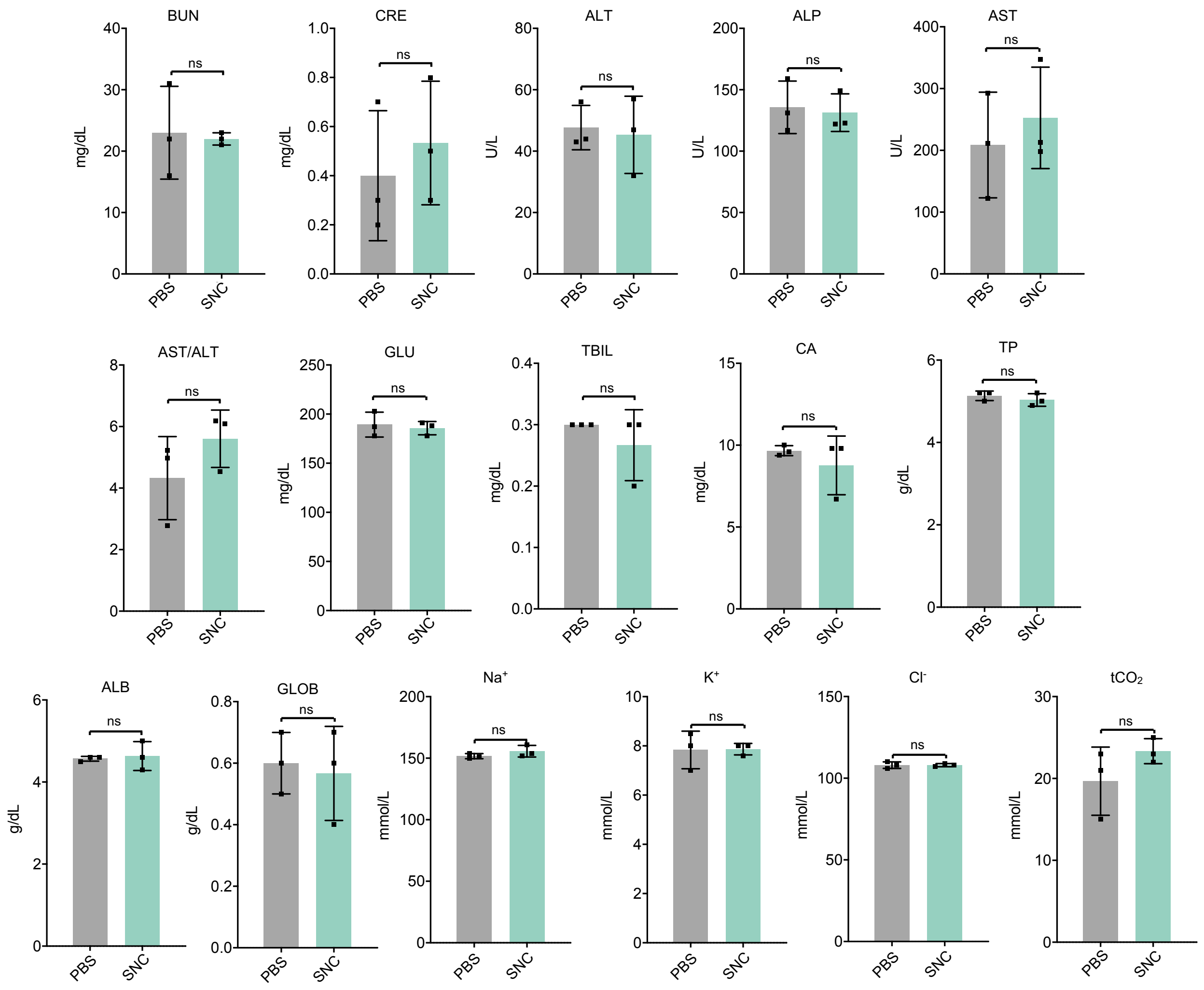


**Figure S20.** Blood biochemical profiles of mice injected with *App*-SNC or PBS (n=3). Fresh whole blood was collected from Ai14 mice on Day 15. ALT, alanine aminotransferase. AST, aspartate aminotransferase. AST/ALT, the ratio of AST/ALT. ALP, alkaline phosphatase. BUN, blood urea nitrogen. CRE, creatinine. TBIL, total bilirubin. GLU, glucose. $Ca^{2+}$, total calcium. TP, total protein. ALB, albumin. GLOB, globulin. $Na^{+}$, sodium. $K^{+}$, potassium. $Cl^{-}$, chloride. tCO2, total carbon dioxide. Data are presented as mean ± s.d. (n=3). Statistical significance was calculated with PBS-injected as the control via one-way ANOVA with Tukey's post hoc test. ns, not significant.

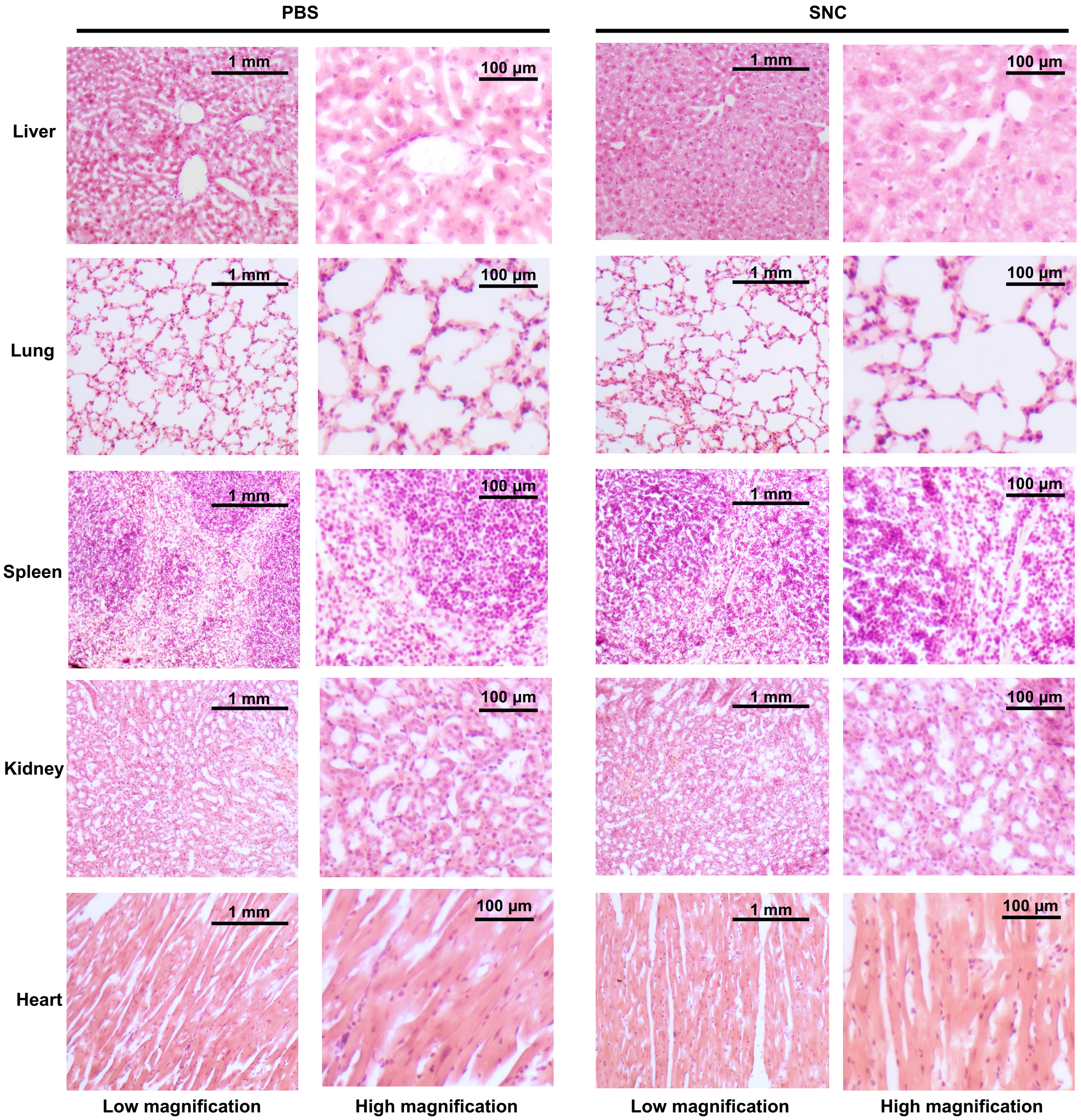


**Figure S21.** H&E staining of major organs of mice received an intravenous injection of PBS and *App*-SNC, respectively. No histological change was observed in different brain regions.

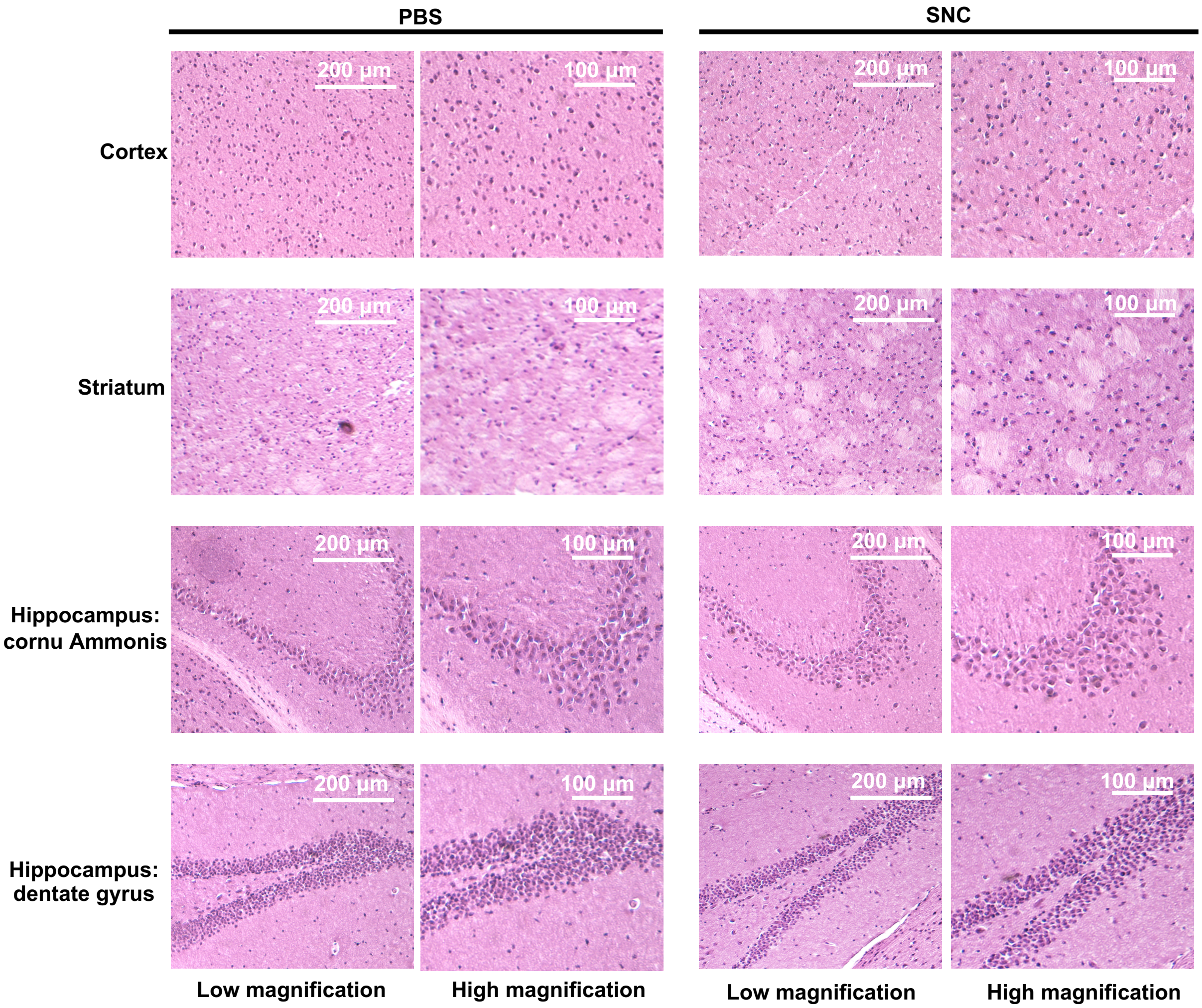


**Figure S22.** H&E staining of mouse brains received an intravenous injection of PBS and *App*-SNC, respectively. No histological change was observed in different brain regions.

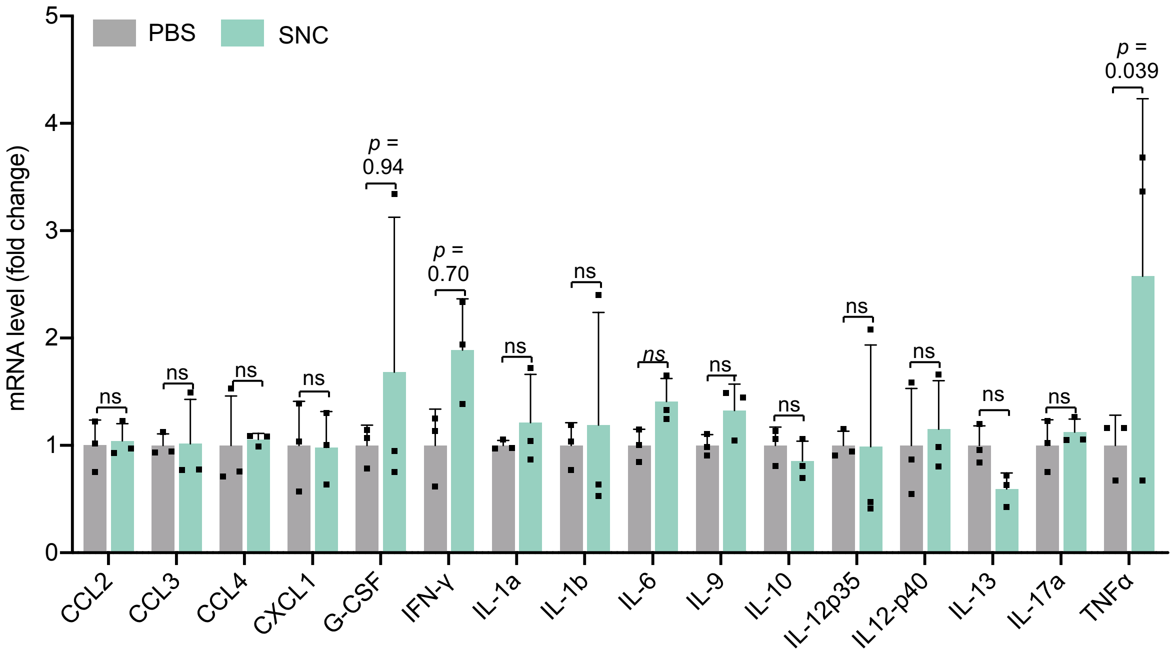


**Figure S23.** Immunogenicity of intravenously injected *App*-SNC in mouse brain. Expression levels of cytokines were quantified by RT-PCR (n = 3). Data are presented as mean ± SD. Statistical significance was calculated with PBS-injected as the control via two-way ANOVA with Tukey's post hoc test. ns, not significant.

**Supplementary Tables**

**Table S1.** sgRNA protospacers.

| Gene | Protospacer |
|---|---|
| *GFP* | 5’-GCACGGGCAGCTTGCCGG-3’ |
| Ai14 (targeting *SV40 polyA*) | 5’-AAGTAAAACCTCTACAAATG-3’ |
| *App* | 5’-ATCCATCCATCATGGCGTGG-3’ |
| *Th* | 5’-CCCCAAGGTTCATTGGACGG-3’ |

**Table S2**. Summary of primary antibodies used for IFCM, CLSM and WB.

| Antibody | Source | Dilution factor | | |
|---|---|---|---|---|
| | | CLSM | IFCM | WB |
| Mouse anti-GAPDH | Thermo Fisher, MA5-15738 | - | - | 1:5,000 |
| Rabbit anti-RFP | Abcam, ab152123 | 1:1,000 | 1:1,000 | - |
| Mouse anti-NeuN | Abcam, ab104224 | 1:1,000 | 1:1,000 | - |
| Mouse anti-GFAP | Abcam, ab190288 | 1:1,000 | 1:1,000 | - |
| Rat anti-CD31 | Abcam, ab56299 | 1:500 | 1:250 | - |
| Rabbit anti-APP (Y188) | Abcam, ab32136 | 1:250 | - | 1:1,000 |
| Rabbit anti-TH | Abcam, ab137869 | 1:250 | - | 1:5,000 |
| Rat anti-EpCAM | Santa Cruz, sc-53532 | 1:100 | - | - |

**Table S3**. Summary of secondary antibodies used for IFCM, CLSM and WB.

| Antibody | Source | Dilution factor | | |
|---|---|---|---|---|
| | | CLSM | IFCM | WB |
| Anti-Rabbit IgG H&L (Alexa Fluor® 594) | Abcam, ab150080 | 1:1,000 | 1:1,000 | - |
| Anti-Mouse IgG H&L (Alexa Fluor® 488) | Abcam, ab150113 | 1:1,000 | 1:1,000 | - |
| Anti-Rat IgG H&L (Alexa Fluor® 647) | Abcam, ab150155 | 1:1,000 | 1:500 | - |
| Anti-Mouse IgG (IRDye® 800CW) | LI-COR, 926-32210 | - | - | 1:5,000 |
| Anti-Rabbit IgG (IRDye® 680RD) | LI-COR, 926-68071 | - | - | 1:5,000 |

**Table S4**. Sequences of primers for PCR.

| Gene | Forward primer (5' to 3') | Reverse primer (5' to 3') |
|---|---|---|
| *App* | TGTCATAGCAACCGTGATTGT | TGCCCTGAGTACCACAGA |
| *Th* | CACAGCCTCCAATGGGTT | TGTTAGTCCTCCACTCCTACAT |
| *CCL2* | GCTACAAGAGGATCACCAGCAG | GTCTGGACCCATTCCTTCTTGG |
| *CCL3* | ACTGCCTGCTGCTTCTCCTACA | ATGACACCTGGCTGGGAGCAAA |
| *CCL4* | ACCCTCCCACTTCCTGCTGTTT | CTGTCTGCCTCTTTTGGTCAGG |
| *CXCL1* | TCCAGAGCTTGAAGGTGTTGCC | AACCAAGGGAGCTTCAGGGTCA |
| *G-CSF* | TCCAGGAGAAGCTGGTGAGTGA | CGCTATGGAGTTGGCTCAAGCA |
| *IFN-γ* | CAGCAACAGCAAGGCGAAAAAGG | TTTCCGCTTCCTGAGGCTGGAT |
| *IL-1α* | CGAAGACTACAGTTCTGCCATT | GACGTTTCAGAGGTTCTCAGAG |
| *IL-1β* | TGGACCTTCCAGGATGAGGACA | GTTCATCTCGGAGCCTGTAGTG |
| *IL-9* | TCCACCGTCAAAATGCAGCTGC | CCGATGGAAAACAGGCAAGAGTC |
| *IL-10* | CGGGAAGACAATAACTGCACCC | CGGTTAGCAGTATGTTGTCCAGC |
| *IL-12p35* | ACGAGAGTTGCCTGGCTACTAG | CCTCATAGATGCTACCAAGGCAC |
| *IL-12p40* | TTGAACTGGCGTTGGAAGCACG | CCACCTGTGAGTTCTTCAAAGGC |
| *IL-13* | AACGGCAGCATGGTATGGAGTG | TGGGTCCTGTAGATGGCATTGC |
| *IL-17a* | GAAGCTCAGTGCCGCCA | TTCATGTGGTGGTCCAGCTTT |
| *TNF-α* | GGTGCCTATGTCTCAGCCTCTT | GCCATAGAACTGATGAGAGGGAG |
| *GAPDH* | TGAGGCCGGTGCTGAGTATGTCG | CCACAGTCTTCTGGGTGGCAGTG |
